# CONNECTOR, fitting and clustering of longitudinal data to reveal a new risk stratification system

**DOI:** 10.1101/2022.08.08.503120

**Authors:** Simone Pernice, Roberta Sirovich, Elena Grassi, Marco Viviani, Martina Ferri, Francesco Sassi, Luca Alessandrì, Dora Tortarolo, Raffaele A. Calogero, Livio Trusolino, Andrea Bertotti, Marco Beccuti, Martina Olivero, Francesca Cordero

## Abstract

The transition from the evaluation of a single time point to the examination of the entire dynamic evolution of a system is possible only in the presence of the proper framework. The strong variability of dynamic evolution makes the definition of an explanatory procedure for data fitting and data clustering challenging. Here we present CONNECTOR, a data-driven framework able to analyze and inspect longitudinal data in a straightforward and revealing way. When used to analyze tumor growth kinetics over time in 1599 patient-derived xenograft (PDX) growth curves from ovarian and colorectal cancers, CONNECTOR allowed the aggregation of time-series data through an unsupervised approach in informative clusters. Through the lens of a new perspective of mechanism interpretation, CONNECTOR shed light onto novel model aggregations and identified unanticipated molecular associations with response to clinically approved therapies.

## 1 INTRODUCTION

In the biological and medical fields, longitudinal data are valuable to explore the evolution of a given event and are expected to have higher predictive power than cross-sectional studies, in which variables are collected at one time point only across a sample population. The investigation of the evolution of a system delivers useful insights into (i) how the measurements change over time within samples; (ii) the time span of relevant events; and (iii) how time evolution is associated with clinical surveillance.

Longitudinal data come in many forms. However, their main characteristic is that they consist of portions of functions or curves, with quantities observed as they evolve through time. In the modeling of temporal data, the state of the art of mathematical methodologies can be classified into two types of approaches: statistical models, in which no biological mechanisms are specified, and mechanistic models, in which all relations are specified (Kendall *et al*., 1999). The statistical approach makes few assumptions (hence, it tends to be biologically naïve) and provides no information about the underlying mechanisms of the system under study. Conversely, being based on a theoretical framework imposed by the modelers, mechanistic models are more suited to yield biologically-aware knowledge. However, due to the complexity of biological systems, it is difficult for mechanistic models to include the intricate relationships that connect all the factors underlying temporal dynamics.

In this context, we propose a workflow based on methods for Functional Data Analysis (Ferraty and Vieu, 2006; Ramsay and Silverman, 2005). The fundamental aims of FDA are those of classical statistics for simple points in a general but finite dimension. However, the classical methods developed for finite dimension and independent observations cannot be directly applied to infinite dimensional data such as functions or curves. Indeed, being functional data, the sampled variables are strongly correlated and the problem becomes ill-conditioned in the context of multi-variate linear models. By analyzing data that vary over time, FDA statistics provide the analytical ground to interpret longitudinal data.

Using FDA methods, here we present CONNECTOR, a data-driven framework to analyze and inspect longitudinal data with the aim to offer a new perspective of mechanism interpretation. CONNECTOR provides several graphical visualizations, supporting users alongside all the analytical steps and required parameters optimizations. Moreover, we distribute a Docker image to guarantee full reproducibility of the presented analyses and ease of use.

To illustrate the effectiveness of this computational analysis workflow, we have leveraged cancer growth data. The collection of cancer growth data increased constantly in the last years. These data, retrieved from different types of biological material, i.e. cancer cell lines (Sharma *et al*., 2010), patient-derived xenografts (PDXs) and organoids (Hidalgo *et al*., 2014; Rizzo *et al*., 2021) are used as pre-clinical models to investigate the mechanisms underlying cancer progression and to identify effective treatments for specific patients’ subsets. In the case of PDXs, a common approach for monitoring tumor growth kinetics consists of evaluating average percentages of tumor volume variations between two time points of interest, hence relying on repetition of the measurements that are presumed independent. The analysis of variance (ANOVA) or t-tests for such values are computed in order to evaluate the punctual time effect among all study arms. In some cases, a categorical system in which the average percentages are classified into specific clinical categories is implemented. In (Oberg *et al*., 2021) a linear mixed effects regression model (with eventually quadratic or cubic terms added) was used to fit and compare the tumor area, after natural logarithm transform. A broad class of mechanistic models can be used for fitting (Kareva I, 2018) data. However, there is no global consensus or biological/clinical evidence on the optimal model that is expected to best fit the growth data (Sarapata and De Pillis, 2014; Benzekry *et al*., 2014).

All previous approaches revolved around a knowledge-based reduction of the systems convolution. CONNECTOR is presented here as a data-driven de-convolution methodology able to fit and cluster temporal data with an accuracy that highlights differences in the dynamics. The CONNECTOR clusters reflected molecular features rooted in the biology of the systems studied. To the best of our knowledge, no other available tools support users without advanced expertise in the statistical analysis of complex temporal data in such an informative and interpretative manner as our proposed software.

CONNECTOR was assessed using tumor growth data from PDX models and resulted in the generation of the CONNECTOR Tumor Growth Classes (CTGC). We first analyzed PDX lines from 36 models generated from chemotherapy-naive high-grade serous epithelial ovarian cancer. CONNECTOR separated the PDX models based on the extent of intra-tumor heterogeneity, which was reflected in the variability of response of individual PDXs to three treatment regimens. Then, we deployed CONNECTOR to inspect the evolution of more than 1500 growth curves of PDXs from metastatic colorectal cancer treated with the clinically approved anti-EGFR monoclonal antibody cetuximab. The clusters generated by CONNECTOR allowed us to identify a subset of cetuximab-resistant tumors associated with previously unrecognized molecular and phenotypic features.

## 2 RESULTS

### Overview of the CONNECTOR framework

CONNECTOR is a tool for the unsupervised analysis of longitudinal data, that is, it can process any sample consisting of measurements collected sequentially over time. CONNECTOR is built on the model-based approach for clustering functional data presented in (James and Sugar, 2003), which is particularly effective when observations are sparse and irregularly spaced, as growth curves usually are.

Hereafter an overview of the software is illustrated (Figure 1), while the full description is reported in the Materials and Methods section. Any collection of observations recorded at several different time points, describing a system evolution, is accepted as a longitudinal data set. These data, together with the description of each sample through a set of relevant features, are the input data sets of CONNECTOR.

**FIGURE 1.**
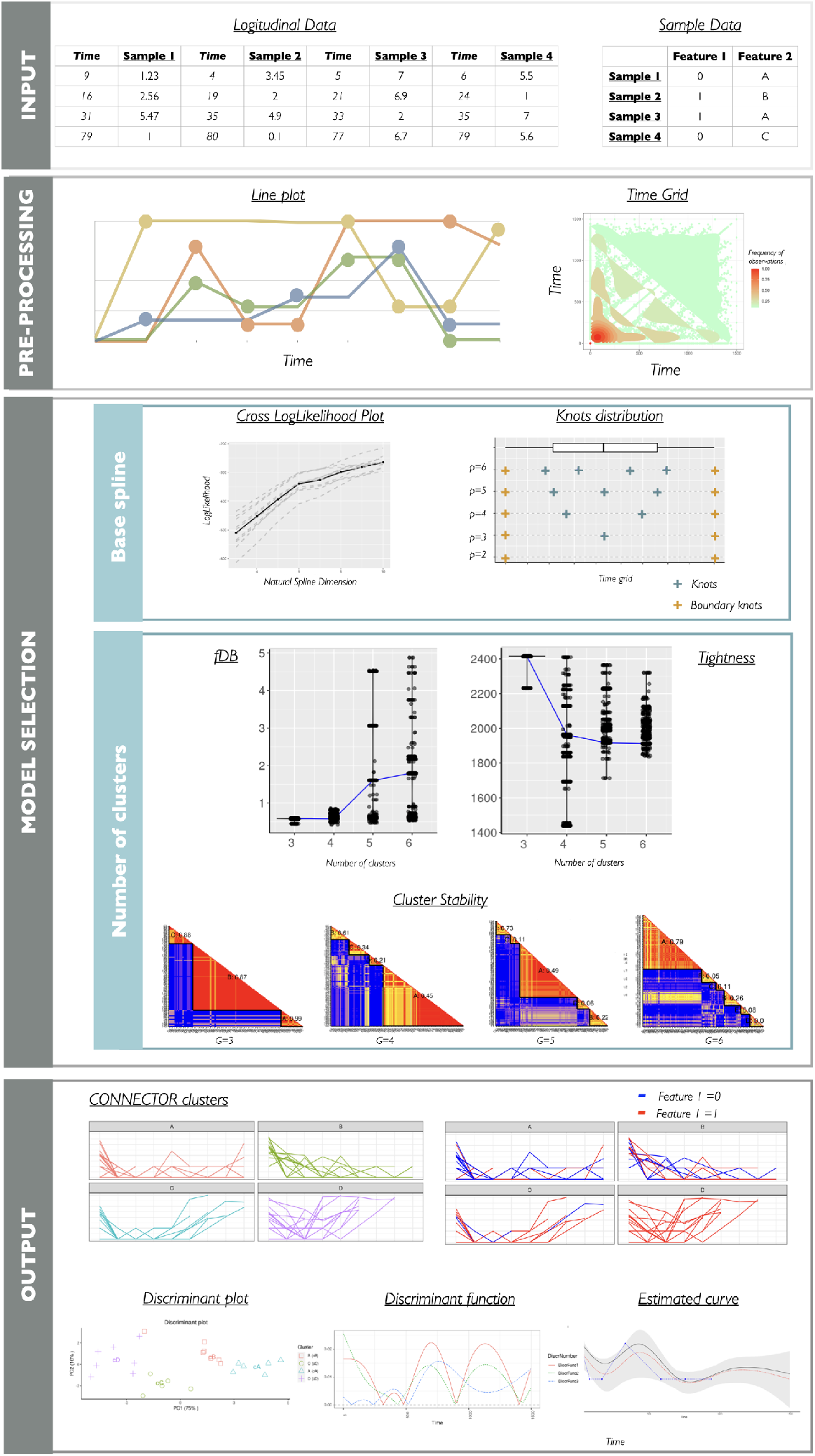
The framework pipeline of the CONNECTOR package. The four main stages of the data processing are illustrated together with the supporting plots returned by CONNECTOR. The **input** data are the sampled curves, which can be also associated to annotation features. Data are **pre-processed** and curves are plotted as lines with dots corresponding to the sampled time points. The heatmap of the full time grid is also provided. The **model selection** is supported with the cross validated loglikelihoods and the knots positions for the choice of the dimension of the spline basis, and with the functional Davies and Bouldin (fDB, see eq. (5)) violin plots, the total tightness (see eq. (4)) violin plots and the stability matrices for the choice of the number of clusters. The **output** of the process is illustrated with the plots of the clustered curves.

The CONNECTOR framework is based on the following steps:

- The *pre-processing step* consists of a visualization of the longitudinal data by a line plot and a time grid heat map helping in the inspection of the sparsity of the time points.
- Then, by means of the FDA analysis, the sampled curves are processed by CONNECTOR with a functional clustering algorithm based on a mixed effect model. The curves are modeled using a finite set of functions (natural cubic splines) with random effects term on the coefficients. The model is fitted and the unknown cluster membership is estimated. This step requires a *model selection phase*, in which several measures are computed that help the user to properly set the two free parameters of the model. The dimension of the spline basis vector is the first free parameter to be set. Two plots, the *cross-log-likelihood* plot and the *knots distribution* plot, are generated for this task. The first is generated by the computation of the cross-log-likelihood exploiting the ten-fold cross-validation method (James *et al*., 2000); the largest stable value of the mean cross-log-likelihood function is attained at the optimal dimension of the spline basis. The second plot visualizes the spline basis knots position for different values of the dimension. The knots divide the time domain into contiguous intervals, and the curves are fitted with separate polynomials in each interval. Hence, this plot allows the user to visualize whether the knots properly split the time domain considering the distribution of the sampled observations. The second free parameter is the number of clusters. The *total tightness* and the *functional Davies-Bouldin (fDB) index* are returned jointly with the stability matrices to support the parameter setting. The violin plots of the cluster dispersion (i.e. tightness) and of the cluster separation (i.e. fDB index) assist the user in the selection of the number of clusters. For the sake of completeness, for different numbers of clusters, the stability matrices are also provided to verify consistency of the samples cluster assignments across different execution runs.
- Once the model selection phase is completed, the *output* of CONNECTOR is composed of several graphical visualizations to easily mine the results, see bottom panel in Figure 1. The dynamics data are plot in clusters, and for each cluster the cluster mean curve is also reported. The curves can be colored using the features associated with each sample as reported in the Sample Data file. The discriminant plot offers a visualization of the sample separation in the CONNECTOR clusters, projected on a plane. Moreover, the discriminant function plot shows the discriminant power of each time point. Finally, the estimated curve, the confidence intervals and the observations are reported for each sample.

#### 2.1 Application of CONNECTOR to PDX models of ovarian cancer

CONNECTOR is extremely flexible in analyzing longitudinal data and aims to tackle a key difficulty in the visualization of high-dimensional clustering data. To show the basic use of CONNECTOR, we analyzed the spontaneous tumor growth patterns of PDX lines derived from the propagation of one chemotherapy-naïve high-grade serous epithelial ovarian cancer (HGS-EOC), which had been passaged for five generations until production of 21 xenografts (Erriquez *et al*., 2016).

The progression curves of the PDX lines are reported in Figure 2A. The time grid suggests that until 70 days the frequency of the observations is larger than 0.75 and it drops for larger times, so we truncated the curves at day 70. The model selection suggests that the optimal parameters set is dimension of the spline basis equals to 3 and number of clusters equals to 4 (CTGC-A, CTGC-B, CTGC-C, CTGC-D). This combination of the parameters led to an average value of the cluster stability equal to 0.8. The detailed CONNECTOR analysis is reported in Figure S1. Figure 2A shows the four CTGCs in which it is possible to appreciate four different progression patterns.

**FIGURE 2.**
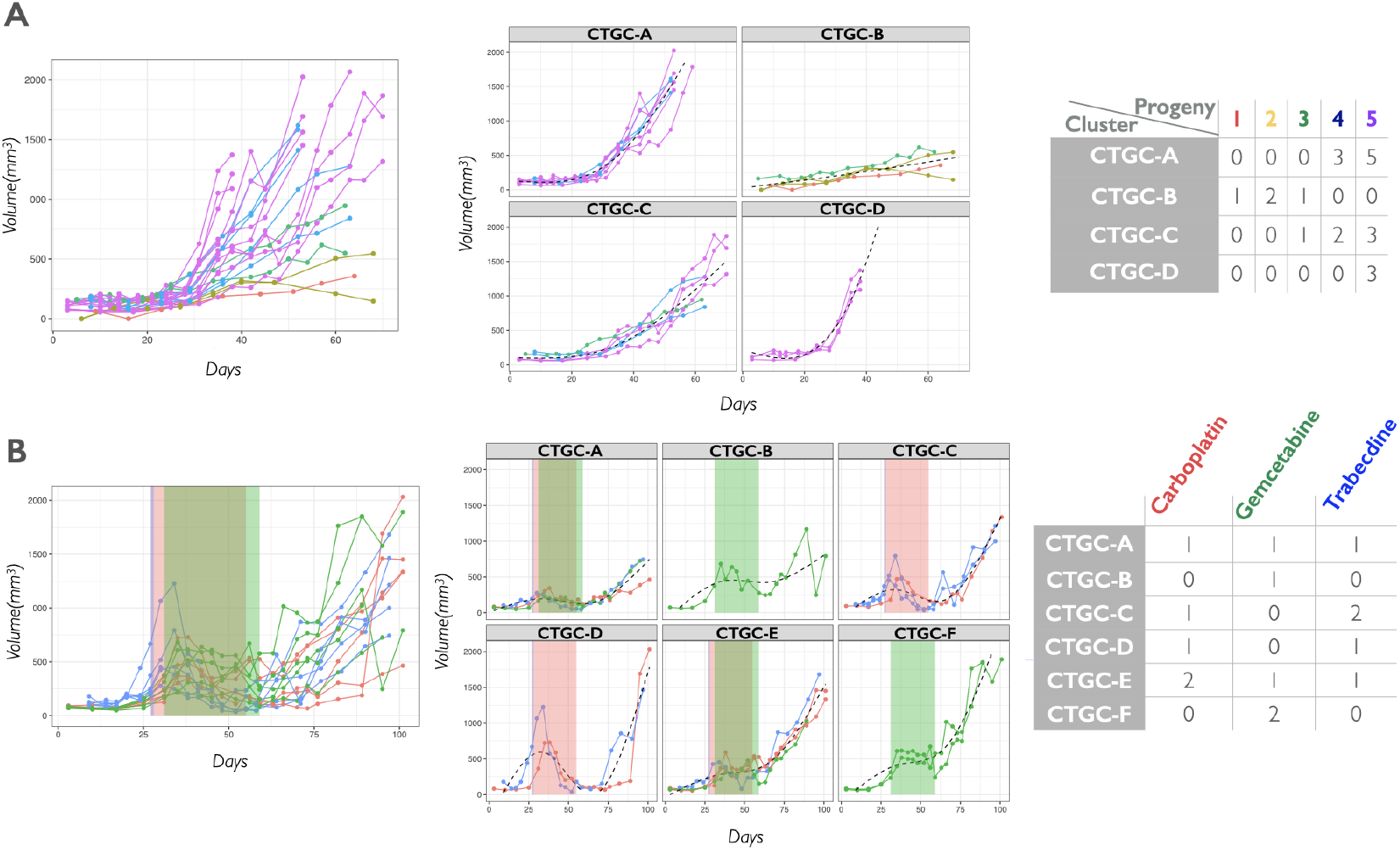
HGS-EOC PDX lines analysis. **A) Untreated PDX lines.** The curves are grouped in four clusters (CTGC-A, CTGC-B, CTGC-C, CTGC-D) and the counts of the different progenies in the different clusters is reported. **B) Treated PDX lines.** The shaded boxes (red, green and blue) highlight the treatments (carboplatin, gemcitabine and trabectedin) time windows (four weeks, four weeks, and once injection - straight blue line at day 28). The curves are grouped in six clusters (CTGC-A, CTGC-B, CTGC-C, CTGC-D, CTGC-E, CTGC-F); the counts of the PDX lines for each treatment in the CTGCs are reported.

Notably, the cluster aggregation reflected a generations separation, likely due to intra-tumor heterogeneity. Indeed, initial engraftment is expected to impose a selection bottleneck, and subsequent propagations exacerbate clonal divergence owing to repeated sampling bias and genomic evolution. During the growth of each PDX line, functional outliers came out, that is, single PDXs emerged that were characterized by increased growth rates. For example, the PDX lines in CTGC-D reached 1000*mm*^3^ of volume in 40 days with respect to the models in CTGC-A or CTGC-C, which reached the same volume in 60 and 70 days, respectively. These outliers might represent the expansion of clonal subpopulations with a fitness advantage. Moreover, the discriminant function returned by CONNECTOR highlights the strong discriminant power of the first 15 days of observation, which can be ascribed to the intra-tumor heterogeneity, see Figure S1.

To establish the versatility of CONNECTOR in analyzing different dynamics, we investigated the tumour growth rates of 15 PDX lines propagated from the same chemotherapy-naive HGS-EOC sample presented above, which were exposed to carboplatin (n=5), gemcitabine (n=5) or trabectedin (n=5). Carboplatin and gemcitabine were administered twice weekly for 4 weeks via intraperitoneal injection; trabectedin eas only once administered through the tail vein. The optimal model selected indicates the dimension of the spline basis equals to 4 and the number of clusters equals to 6 (CTGC-A, CTGC-B, CTGC-C, CTGC-D, CTGC-E, CTGC-F); considering this setting the cluster stability is 0.9. The detailed CONNECTOR analysis is reported in Figure S2. In this cohort of PDXs, the response to drugs of individual models was uneven; some models showed significant reduction, while other models proved to be resistant, Figure 2B. Notably, the discriminant function reveals that the time with the highest discriminatory power corresponds to day 35. Indeed, around that time, the curves start to be separated as a consequence of the chemotherapy treatments effects. In detail, in CTGC-A, CTGC-B and CTGC-F, the tumors remained stable during the treatment time window, while in CTGC-C and CTGC-D a clear increment, followed by a reduction of the tumor mass, was appreciable. This variable distribution of responses can be explained, again, with a high degree of intra-tumor heterogeneity in the original tumor from which the different PDX lines were propagated; indeed, genetic deviation is expected to influence response to therapy (Schmitt *et al*., 2016).

#### 2.2 Application of CONNECTOR to PDX models of metastatic colorectal cancer

To show the full potential of CONNECTOR in a complex use-case, we analyzed a large dataset of tumor growth curves of patient-derived xenografts (PDXs) from mice treated with cetuximab, an anti-EGFR antibody that is approved for clinical use in patients with RAS/RAF wild-type metastatic colorectal cancer (mCRC). Results from these xenotrials were obtained from a continuously expanding collection of ~400 mCRC PDXs, part of which had been used in previous studies (Bertotti *et al*., 2011, 2015; Zanella *et al*., 2015; Isella *et al*., 2017; Lupo *et al*., 2020). tumor volumes were measured weekly after tumor implantation and over the course of treatment. To obtain comparable data on tumor regression or growth under treatment, factoring away different engraftment efficiencies, therapy was started when the average tumor volume in the cohort expanded from an individual PDX reached 400 *mm*^3^ and was administered for three weeks.

In previous projects we categorized PDX response to cetuximab based on parameters that are loosely inspired to the RECIST clinical criteria (Ko *et al*., 2021; Bertotti *et al*., 2011). This classification includes three classes, based on the average tumor volume variation at endpoint compared with tumor volume at baseline (the day before treatment initiation): partial responses (PR) are defined as tumors that regress by 50% or more during treatment; progressive diseases (PD) are defined as tumors that increase their volume by 35% or more, despite treatment; tumor volume changes above PR or below PD thresholds are defined as stable diseases (SD). To allow a direct comparison of CONNECTOR’s clustering results with our historical annotation, we decided to analyze, for each tumor, the available measurements between the day before treatment initiation and the following 3 weeks. The selected dataset was extracted from the laboratory LIMS (Baralis *et al*., 2012) and comprises measurements from 1563 individual mice, collected from 2012 to 2020 and representing 173 original engraftments from parental tumors in patients.

The PDX curves were analyzed by CONNECTOR to achieve a widespread overview of the distribution of the individual PDX models after drug administration. When running CONNECTOR, based on the results from the model selection phase, the optimal value of the base spline was 4. To perform a through analysis we evaluated fDB and tightness for a broad range of clusters numbers, starting with three, that corresponds to the number of clinical response classes. We chose to compare the results obtained with 3, 4 and 5 clusters, as the fDB index worsen for larger values, see Figure S3A.

We studied how the PDX growth curves are distributed *intra-parameter setting*, among the number of CTGCs selected for each run, and *inter-parameter setting*, exploring how the curves move among the CTGCs along the three runs (Figure 3A). We observed that by increasing the number of CTGCs, the curves that were separated were mainly those characterized by a sudden volume increase. Based on such analysis, the dataset was optimally described by three primary classes, namely CTGC-A, -B and -C, leading to an average stability matrix of 1. We then assessed whether a higher resolution could be achieved by segregating the larger CTGCs into subclasses. We thus processed again, as independent datasets, the CTGCs composed by more than 200 curves (namely CTGC-A and -B, see Figure S3B and Figure S3C, respectively). Through this, we obtained a final number of 9 CTGCs - where CTGC-A and B are further split in Aa, Ab, Ac, Ad, Ae and Ba, Bb, Bc - which are represented in Figure 3B.

**FIGURE 3.**
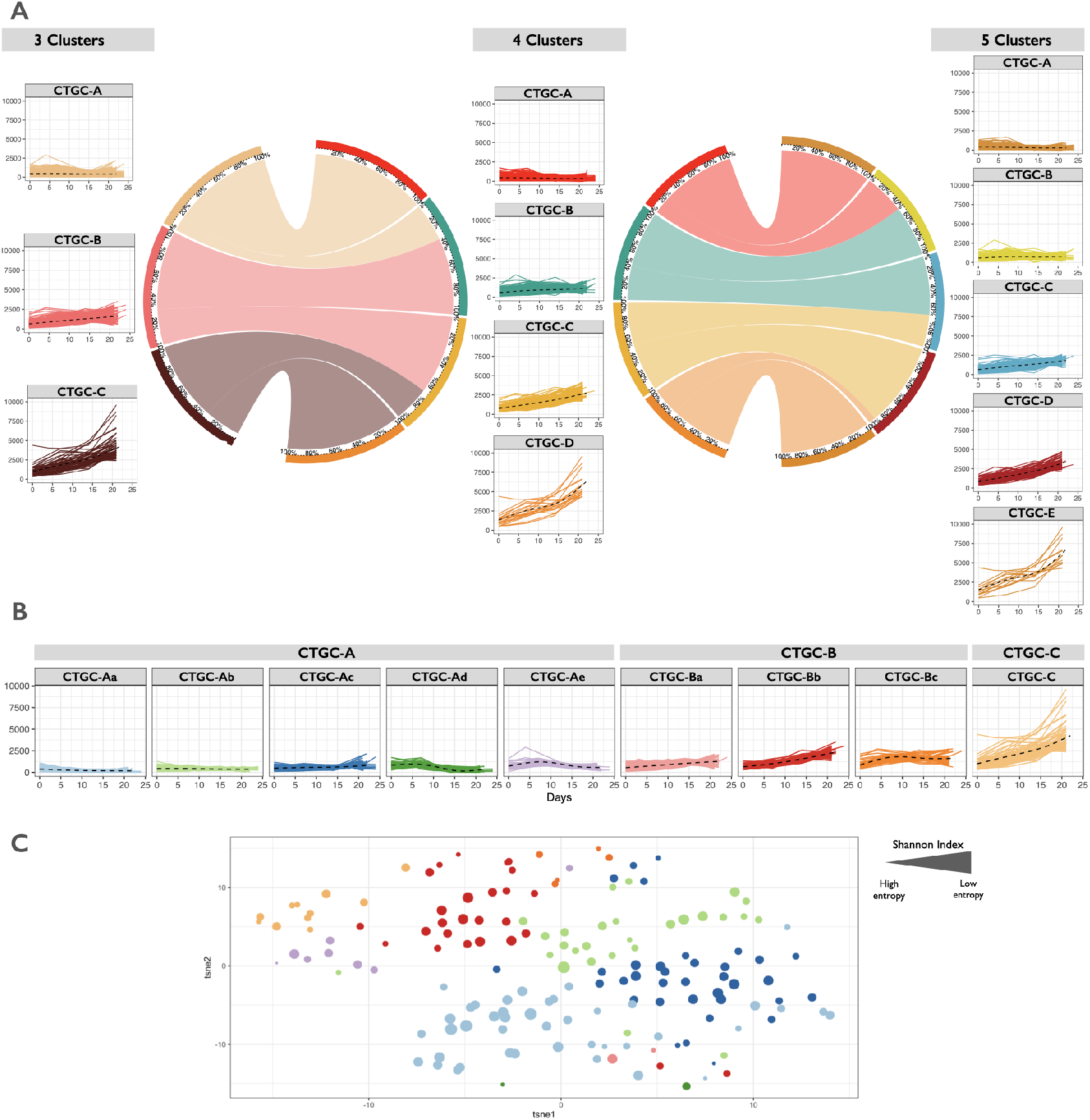
CONNECTOR results. **A) CONNECTOR tumor growth classes** for three different choices of the number of clusters: three, four and five. The circos plots show the repositioning of the curves as the number of CTGCs changes. **B) CONNECTOR tumor growth classes with twofold clustering.** The nine boxes result from a first run with number of clusters equal 3 followed by second runs on the CTGC-A (with 5 sub-classes) and on the CTGC-B (with 3 sub-classes). **C) t-SNE visualization of the CTGCs induced on models.** The color of the dots match the color of the assigned CTGC, represented in panel B. The dimension of the dots is inversely proportional to the Shannon Index calculated on the distribution of the curves of the same model across CTGCs (large dots - small entropy).

We assigned each PDX model to a specific CTGC, by means of a Naïve Bayesian classificator, see Section 4. By calculating the Shannon index representing the consistency of the curves obtained from the same model, we observed that in most cases there is high agreement among the CTGCs assigned to different curves of the same parental tumor (mean = 1.54, s.d. = 0.52). For the sake of clarity, an additional graphical visualization, based on the t-distributed stochastic neighbour embedding (t-SNE) is proposed. The plot is included in Figure 3C, where distinctly isolated CTGCs can be appreciated, in good agreement with the CONNECTOR CTGCs.

#### 2.3 CONNECTOR clusters reveal a subgroup of non-responder tumors that express high levels of stratified epithelium keratins

Different CTGCs were evaluated with respect to our three-class cetuximab response annotation and the molecular characteristics of the classified tumors (Figure 4A).

**FIGURE 4.**
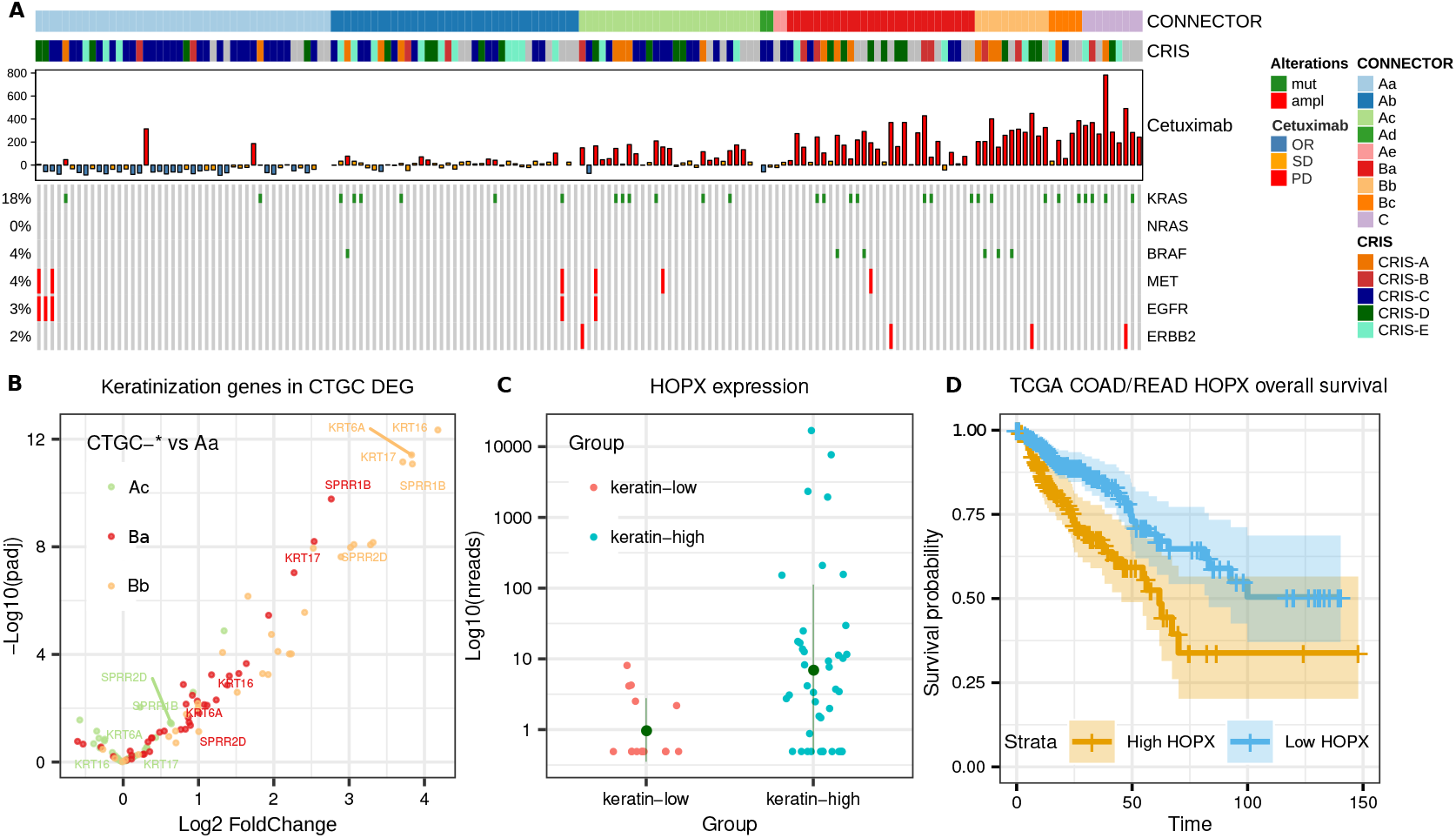
CONNECTOR clusters molecular annotation and transcriptomic analyses. **A) Molecular and phenotypic characterization of the CONNECTOR clustered mCRC xenografts:** each sample was annotated according to CRIS subtype, response to cetuximab and somatic alteration known to determine cetuximab resistance (KRAS, NRASand BRAF single nucleotide variations; MET and ERBB2 amplification) or sensitivity (EGFR amplification). **B) Differential expression of genes in the “keratinization” GO in CTGCs**: triple volcano plot showing the magnitude of expression differential (x axis, Log2 FoldChange) and significance (y axis, -Log10 adjusted p value) of “keratinization” genes when comparing CTGCs enriched in PD versus Aa. Only comparisons involving CTGCs enriched in PD with at least 9 total samples are reported for clarity. Bb and, to a smaller extent, Ba have many keratinization genes upregulated, but the same is not true for Ac. The top 5 upregulated genes in Bb are labeled (SPRR is a family of small proline rich proteins induced during differentiation of keratinocytes). **C) Expression levels of HOPX in keratin-high and keratin-low samples:** the y axis shows DESeq2 corrected counts in logarithmic scale. **D) Survival analysis of TCGA colorectal patients stratified by HOPX expression levels:** overall survival time is in month, the high group comprises 134 patients, the low 237. Log rank p value 8.6 × 10^-3^.

As expected, CTGCs were associated with RECIST-like response classes. Indeed, PR tumors were enriched in CTGC-Aa (Fisher p value 2.4 × 10^-13^) and SD tumors were enriched in CTGC-Ab (Fisher p value 8.0 × 10^-8^). Notably, CONNECTOR segregated PD tumors in multiple clusters (CTGC-Ac, p value 5.5 × 10^-2^; CTGC-Ba, p value 1.9 × 10^-4^; CTGC-Bb 7.5 × 10^-5^, p value; CTGC-Bc, p value 1.7 × 10^-1^; CTGC-C, p value 4.6 ×10 ^-4^), indicating that CONNECTOR can distinguish different growth patterns within the PD standard response category.

The analysis of CTGC associations with the transcriptional subtypes identified by a PDX-based, cancer cell-intrinsic (CRIS) classifier (Isella *et al*., 2017) showed interactions between the growth patterns detected by CONNECTOR and specific biological traits. In line with the observation that cetuximab-responsive tumors are enriched in the CRIS-C subtype (Isella *et al*., 2017), CRIS-C tumors were associated with the CTGC-Aa cluster. However, the enrichment of CRIS-C tumors within CTGC-Aa (Odds Ratio CRIS C/Aa 4.63) was stronger than that observed when considering all responders (Odds Ratio CRIS-C/PR 3.38). This suggests that CONNECTOR is able to recognize the specific subset of responders sustained by the CRIS-C phenotype.

The finding that CONNECTOR clusters accurately captured defined molecular features prompted us to investigate whether the diversification of cetuximab-resistant cases in multiple CTGCs could relate to different biological substrates of resistance. To test this hypothesis, we first assessed the distribution of the genetic variants that are known to cause resistance to cetuximab in colorectal tumors (Bertotti *et al*., 2015). As expected, all such variants were depleted from CTGC-Aa (Fisher p value 1.3 × 10^-2^) and, albeit to a non-significant extent, from CTGC-Ab (Fisher p value 0.22); conversely, resistance variants were overall enriched in all the CTGC-B clusters and CTGC-C cluster. Importantly, we did not observe an enrichment for specific resistance mutations in any of the PD-associated CTGCs, suggesting that the tumor growth patterns recognized by CONNECTOR in non-responders were not driven by specific resistance genotypes.

To further explore the functional characteristics of the different PD-enriched CTGCs, we mined a set of RNA-seq data that were generated in the context of an independent study performed in the same cohort (Perron et al, in preparation). Analysis of differentially expressed genes (DEGs) between all PD-enriched CTGCs and CTGC-Aa, used as a common reference, uncovered a very specific GO functional enrichment related to stratified epithelial differentiation and keratinization, which was guided by keratin-encoding genes significantly upregulated in CTGC-Ba and CTGC-Bb, but not in CTGC-Ac (Figure 4B, Table S1 and Figure S4E-F). This was not simply driven by the enrichment of non-responder tumors in CTGC-Ba and CTGC-Bb. Indeed, the same functional enrichments for CTGC-Ba and CTGC-Bb tumors versus CTGC-Ac tumors were observed also when limiting the analysis to PD tumors only (see Table S2 and Figure S4A-B-C). Hence, CTGC unveiled two distinct phenotypes associated with cetuximab resistance, one of which (shared by CTGC-Ba and CTGC-Bb) is characterized by markers of keratinized stratified epithelia.

To validate this observation, we stratified the full cohort of cetuximab-resistant PDXs for which RNAseq data were available (n = 140) based on the expression of 10 keratin genes that were robustly upregulated in CTGC-Ba or CTGC-Bb and associated to “keratinization” according to GO annotations (details in Section 4). By applying thresholds derived from expression quartiles in CTGC-Ac non-responsive tumors, we identified 42 keratin-high samples and 15 keratin-low samples. H&E sections of 4 keratin-low and 4 keratin-high PDXs were subjected to blinded histopathological evaluation, which confirmed the existence of two clearly distinct subpopulations (see Figure S5). The first included tumors displaying histological features that resembled the glandular organization typical of the large intestine (presence of adenomorphic structures with evident luminal spaces, mild to moderate tumor cell pleomorphism, columnar cells with nuclear polarization). tumors pertaining to the second group were less differentiated, with round or columnar pleomorphic cells that formed solid fields and rare luminal spaces. Intriguingly, in three of the latter four tumors we observed a variable degree of squamous differentiation, with evidence of keratin pearls in one of them. Indeed, after unblinding, three of the selected keratin-high tumors were found to belong to the latter group, confirming that CTGC-B gene expression traits were likely associated with the acquisition of histological traits typical of squamous epithelia.

DEG analyses comparing keratin-high and keratin-low tumors confirmed the GO functional enrichments previously observed in CTGC-B tumors (see Table S3 and Figure S6). Furthermore, novel associations to terms related to cell motility, wound healing and angiogenesis emerged. This may indicate that the keratin-high subset is endowed with more aggressive and invasive properties with respect to keratin-low tumors. Interestingly, we found a robust enrichment for CRIS-B (fisher p value 2.09 × 10^-6^) among keratin-high tumors. CRIS-B was previously reported as an aggressive transcriptional subtype associated with poor prognosis (Isella *et al*., 2017) and undefined etiology. Our data may indicate that the CRIS-B phenotype is sustained by an aberrant differentiation pattern towards epithelial cornification. In agreement, some of the keratins typical of CTGC-B (KRT17, KRT6A, KRT80) are also marker genes for CRIS-B, further reinforcing the link between the two classes.

The keratinocytic transdifferentiation of CTGC-B tumors may be driven by activation of a stemness-related pathway sustained by the transcription factor HOPX, which is indeed upregulated in our keratin-high subgroup (Log fold change 3.1, adjusted p value 4.23 × 10^-6^, Figure 4C). HOPX is a known regulator of differentiation lineages (Liu and Zhang, 2020) in various tissues, and sustains the reservoir of intestinal stem cells that can originate all intestinal epithelial compartments (Takeda *et al*., 2011). Although the role of HOPX in CRC is controversial (Yamashita *et al*., 2013; Dmitrieva-Posocco *et al*., 2022), the assumption that its activation could contribute to the progression of a subset of aggressive tumors is in agreement with the observation that high expression of HOPX is significantly associated with bad prognosis in CRC (overall survival Log Rank p value 8.6 ×10^-3^ in the TCGA colorectal cohort, Figure 4D and (Liu and Zhang, 2020)).

The connection between CRC cornification and HOPX activity is supported by the role of HOPX in the epidermis, where HOPX-positive precursors give rise to a set of keratin 6-positive inner bulge cells of the hair follicle (Takeda *et al*., 2013). This may explain why HOPX activation, besides sustaining the progression of aggressive and invasive tumors, also induces an aberrant transdifferentiation towards keratinized lining epithelium, which ultimately associates with the tumor growth pattern typical of class B CTGC.

## 3 DISCUSSION

The main goal of this study was to develop a tool for exploring longitudinal data with an unsupervised approach. Hence, we worked on a tool for aggregating curves into classes that could suggest directions and hypotheses for the in-depth examination of different datasets.

We reviewed the current literature on FDA and chose to build CONNECTOR based on the functional clustering method proposed in (James and Sugar, 2003). Indeed, depending on the sampling strategy (high frequency or sparse, regular or irregular), different methods have been proved efficient to the clustering task. We were interested in curves that are observed at sparse and irregular times, as usually occurs when data collection is laborious and time limits are constrained by ethical protocols. On these premises, and considering the results illustrated in (Jacques and Preda, 2014), we compared the best reported methods on sparse and irregularly sampled curves. The tests indicated that (James and Sugar, 2003) was the best performing method. The method has been studied to solve the main problems that arise when clustering sparse and irregularly sampled functional data: large number of missing observations on the discretised time grid (sparsity) and different covariances of the coefficients (curves are measured at different time points). Indeed, the method uses basis functions to project each curve onto a finite-dimensional space and considers a random-effect model for the coefficients. This allows to take advantage of all the sampled curves. The model is extremely flexible and computes estimates, confidence intervals and prediction intervals for individual curves.

The performances of the functional clustering model are strongly dependent on the choice of several free parameters. CONNECTOR includes a toolset for choosing each free parameter appropriately. To this goal two new indexes have been introduced – the functional Davies and Bouldin (fDB) index and total tightness. The two indexes, together with the cross-loglikelihood plot, the visualization of the positions of the time knots and the stability matrix of the final clustering, provide the user with all the information needed to choose suitable values of the free parameter.

Oberg and co-workers (Oberg *et al*., 2021) have recently proposed a linear mixed effects regression models for analysis of PDX repeated measures data. The model was able to fit different treatment designs and randomization schemes. However, their model is limited in the prediction of the trajectories since only monomials up to cubic degree are considered to describe the time dependency, after log transform. Conversely, the core of the CONNECTOR package is a general mixed effect model where both fixed-effects term and random-effects term are considered on a cubic spline basis, which makes the model as flexible as needed.

We illustrated the versatility of CONNECTOR starting with the analysis of the spontaneous growth patterns in 21 PDX lines propagated from a single HGS-EOC tumor sample. We observed uneven growth rates in PDX lines derived from the same original tumor. This is consistent with the notion that ovarian cancers show a high degree of intratumor heterogeneity (McPherson *et al*., 2016; Schwarz *et al*., 2015; Castellarin *et al*., 2013), which results in the establishment of genetically different tumor entities during serial propagation. This diversification can also explain the varied responses to treatment observed in 15 PDX lines – again derived from the same original tumor – exposed to different chemotherapeutics.

CONNECTOR allowed us to perform an in-depth study of the growth dynamics of a vast cohort of mCRC PDXs treated with cetuximab, with validation and discovery aspects. First, the reliability of the unsupervised and data-driven identification of CTGCs is supported by the observation that clusters were properly enriched in the three classes by which cetuximab response was previously annotated. Moreover, our analysis adds new insights into how cetuximab-resistant tumors can be stratified according to their molecular diversification. Specifically, CTGC-based categorization allowed us to identify a subset of cetuximab-resistant tumors (CTGC-B) with transcriptional and morphological features of metaplastic differentiation towards cornified epithelia (keratin-high tumors). CTGC-B tumors were also enriched for the CRIS-B transcriptional subtype, which was reported as a poor-prognosis tumor subgroup, composed of highly invasive tumors that are resistant to currently available therapeutic options (Isella *et al*., 2017).

Through CONNECTOR, we identified a previously unnoticed characteristic of such tumors, which may - at least partially - explain their aggressiveness and, at the same time, their keratinized phenotype. A literature search put forward the transcription factor HOPX as a potential regulator of the transdifferentiation process experienced by the keratin-high subgroup of cetuximab-resistant tumors (Takeda *et al*., 2013). HOPX is involved in the modulation of stem cell renewal in both the intestine and the epidermis (Mariotto *et al*., 2016). In the epidermis, HOPX is responsible for the production of keratin 6-positive cells, which contribute to the post-injury regeneration of cornified epithelia by entering proliferation at the wound edge (Moll *et al*., 2008).

HOPX proved to be strongly upregulated in keratin-high, CTGC-B CRC tumors, and we provide evidence that HOPX expression correlates with poor prognosis in CRC patients. On this ground, we speculate that the same program that regulates the regeneration of keratinized epithelia may be aberrantly activated in CRC by HOPX overexpression, thus committing cancer cells towards an aggressive phenotype with traits of epithelial cornification. In this regard, it is worth noting that high expression of keratin 80, one of the markers of the keratin-high subgroup, has been associated with poor prognosis in CRC and with CRIS-B specific phenotypic traits, such as epithelial to mesenchymal transition and cell invasion (Li *et al*., 2018). This finding may have substantial implications. Indeed, if functionally validated, the notion that HOPX is a driver for the maintenance of keratin-high tumors with keratinocytic metaplastic differentiation may pave the way for the development of new therapeutic approaches aimed at interfering with HOPX function. We also note that keratinization did not stand out as a predominant characteristic of CRIS-B in the original study (Isella *et al*., 2017), possibly because the phenotype is specifically associated with the subpopulation of CRIS-B tumors that are resistant to anti-EGFR therapy. This makes CONNECTOR a versatile tool for discerning relevant trends from longitudinal data, which are harder to detect using more classical sources of molecular and phenotypic annotation.

Preclinical cohort studies involving the collection of longitudinal high-dimensional data are being increasingly conducted to evaluate drug activities and to explore disease evolution, and results from such efforts show potential to be translated into experimental trials and clinical practice. Disease monitoring over time is also an already well-established method that is routinely incorporated in several clinical trials. The CONNECTOR framework was designed and implemented as a user-friendly tool to streamline gathering this kind of information, while providing hints to increase interpretability and molecular accuracy.

## 4 MATERIALS AND METHODS

### Functional Data Analysis

#### Functional Clustering Model

The functional clustering method implemented in CONNECTOR is based on the functional clustering model presented in (James and Sugar, 2003). Let us denote as *g_i_*(*t*) the curve of the *i*th selected individual. In practice, we observe *g_i_*(*t*) with measurement errors and only at few discrete time points. Let **Y**_i_, be the vector of observed values of *g_i_*(*t*) at times *t*_*i*_1__,…, *t_i_n_i___*. Then we have

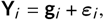

where **g**_*i*_, and *ε_i_*, are the vector of true values and measurement errors at time grid, respectively. As there are only finite number of observations, individual curves are modeled using basis functions, in particular cubic splines. Let

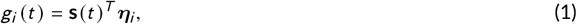

where **s**(*t*) is a *p*–dimensional spline basis vector and **η**_*i*_, is a vector of spline coefficients. The **η**_*i*_’s are treated with a random-effects model. In particular, the spline coefficients are modeled using a Gaussian distribution,

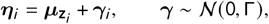

where **z**_*i*_, denotes the unknown cluster membership. Let *G* denote the true number of clusters. Cluster means are furthermore rewritten as

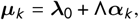

where **λ**_0_ and ***α**_k_* are *p*– and *h*– dimensional vectors, Λ is a (*p, h*) matrix and *h* ≤ min (*p, G* – 1). Thus, *h* represents the dimension of the mean space, allowing a further lower-dimensional representation of the curves with means in a restricted subspace (for *h* < *G* – 1).

With this formulation, the functional clustering model can be written as

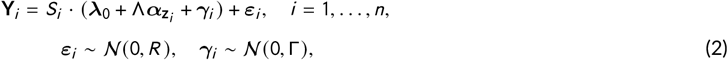

where *S_i_*, = (**s**(*t*_*i*_1__),…,**s**(*t*_*i*_*n_i_*__))^*T*^ is the spline basis matrix for the *i*–th curve. There are many possible forms for *R* and Γ. For now, we use *R* = *σ*^2^*I* and a common Γ for all clusters, as we are interested in sparse datasets, for which the smallest number of parameters is advisable.

The model is fitted following (James and Sugar, 2003) and all the estimated parameters and the predicted cluster membership are returned.

#### Model Selection

Two free parameters have to be properly chosen before fitting: the dimension of the spline basis *p* and the number of clusters to fit *G*.

The dimension of the spline basis can be chosen as the one corresponding to the largest cross-validated likelihood, as proposed in (James *et al*., 2000). CONNECTOR uses a ten-fold crossvalidation, which involves splitting data into 10 roughly equal-sized parts, fitting the model to 9 parts and calculating the loglikelihood on the excluded part. Repeat 10 times and combine loglikelihoods. Notice that, the resulting plot of the mean tested likelihoods versus the dimension of the basis, should be treated as a guide rather than an absolute rule, keeping in mind that working with sparse data pushes to spare parameters. Moreover, as the position of the knots depends on their number, CONNECTOR returns a plot with this information as the parameter *p* varies.

The number of clusters *G* must be chosen. CONNECTOR provides two different plots to properly guide in one of the most difficult problems in cluster analyzis. As in the finite-dimensional case, where data are points in place of curves, we need some proximity measures to validate and compare results of a clustering procedure.

We chose to follow (Ferraty and Vieu, 2006) and rely on the parameterized family of semi-metrics between curves defined as

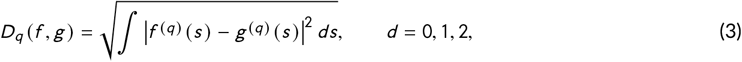

where *f* and *g* are two curves and *f*^(*q*)^ and *g*^(*q*)^ are their *q*th derivatives. Note that for *q* = 0, eq. (3) is the distance induced by the classical *L*^2^–norm. It turns out that *D_q_* can be reliably calculated in our setting where we are interested in proximity measures between each curve in a cluster and the centre-curve of the cluster (tightness of the cluster), as well as proximity measures between each centre-curve and the centre-curves of different clusters (separateness of clusters). Hence we may have *f* being the estimated *i*th curve and *g* being the estimated mean curve of cluster *k*, or *f* and *g* being both mean curves. In any case, *D_q_* can be calculated taking advantage of the spline representation of the estimated curves and mean curves, see eq. (1). Indeed, the computation of successive derivatives, which is numerically sensitive, can be performed by differentiating their analytic form. It remains to compute the integral which can be done by numerical method. As the proper calculation of *D_q_* is a basic condition for the tools shown below, all details are illustrated in the Supporting Information S1.

We define a first quantity to infer the appropriateness of the data partition, which we call *total tightness*. It is the dispersion measure given by

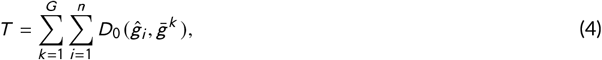

where ĝ, is the estimated *i*-th curve given in eq. (S2) and 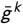 is the center of *k*-th cluster given in eq. (S1), see Supporting Information S2. As the number of clusters increases, the total tightness decreases to zero, the value which is attained when the number of fitted clusters equals the number of sampled curves. In this limiting case, any *k*-th cluster mean curve coincides with an estimated curve and 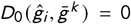 for any *i* and *k*. A proper number of clusters can be inferred as large enough to let the total tightness drop down to relatively little values but as the smallest over which the total tightness does not decrease substantially. Hence, we look for the location of an “elbow” in the plot of the total tightness against the number of clusters.

We define a second index, which is a cluster separation measure. Following (Davies and Bouldin, 1979), we extend the well known Davies-Bouldin (DB) index to the functional setting. Let us call the new index *functional* DB (fDB). It is defined as follows

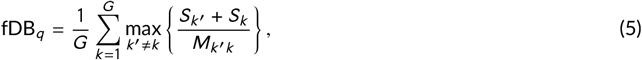

where, for each cluster *k* and *k*′

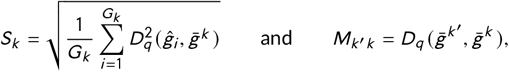

with *G_k_* the number of curves in the *k*-th cluster. The significance of eq. (5) remains unchanged with respect to the finite-dimensional case. It is the average of the similarity measures of each cluster with its most similar cluster. The “best” choice of clusters, then, will be that which minimizes this average similarity. It should be noted that *M_k′k_* is the distance between centroids (mean-curves) of *k*′-th and *k*-th cluster. It serves to weight the sum *S*_*k*_′ + *S_k_*, which is the total standard deviation of the clusters: *k*′-th and *k*-th clusters dispersion is measured compared to their relative distance.

CONNECTOR returns the violin plots of both the total tightness and the fDB index for a given repetition of runs and for different choices of the number of clusters *G*. To support this decision, CONNECTOR returns as well the consensus matrix for the most frequent clustering at each given number of clusters. The plot informs about the stability of the final clustering across different runs. Indeed, each cell of the matrix is colored proportionally to the frequency of the two corresponding curves belonging to the same cluster across different runs. Hence the larger the frequencies are (which corresponds to warmer colors of the cells in the plot), the more stable is the final clustering. Motivated by its meaning, we refer to such matrix as the stability matrix.

The observation of the fDB violin plots, the total tightness violin plots together with the stability matrix enables the user to properly set the free parameter G, which represents the number of clusters to be fitted.

Notice that the functional clustering method allows for a lower dimensional representation of the curves through the parameter *h*. This reduction is often needed as data could be not enough to estimate the large number of parameter of the functional clustering model. CONNECTOR optimizes the choice of the parameter *h* returning the largest value for which the estimation of the parameters is successful with a reasonable, set by the user, frequency. Hence the value of *h* is not chosen directly, but returned by CONNECTOR.

#### Functional Clustering Tools

In (James and Sugar, 2003) three important tools to analyze the clustering are presented: the discriminant plot, the discriminant function and the estimation of the entire curve for each single subject.

The *discriminant plot* is a low dimensional plot of the curves dataset. It helps to visualize the clusters, as each curve is projected in a low dimensional space so that it can be plotted as points. In particular, each curve is represented by its projection onto the h-dimensional space spanned by the means *μ_k_*.

The *discriminant functions* are plots of the weights Λ^T^*S^T^*Σ^-1^, versus time, to apply to each dimension for determining cluster membership. The term *S*Λ is a measure of average separation between clusters and Σ is a measure of their variability. There will be *h* discriminant functions and each curve shows the times with higher discriminatory power, which are the times corresponding to largest absolute (positive or negative) values on the y-axis.

The functional clustering procedure predict unobserved portions of the true curves for each subject. The *estimated curves* are returned by CONNECTOR and plotted with confidence intervals as well. In Supporting Information S3 is reported an exhaustive review on the performance of different procedures with respect to the results obtained by CONNECTOR.

### The CONNECTOR framework

The CONNECTOR software is composed of two main modules which cover the whole analysis, consisting of an R library, called CONNECTOR Package, and a docker image. Docker containerization is utilized to simplify the distribution, use and maintenance of the analysis tools; while the R library provides an easier user interface for which no knowledge of the docker commands is needed.

The framework pipeline is defined by three necessary steps:

1. the data importing and processing to create the R object exploited through the entire analysis,
2. the model selection by identifying the optimal values of the two free parameters of the FCM approach (the basis spline dimension *p* and the number of clusters G),
3. the inspection and visualization of the obtained clusters.

In this contest, the CONNECTOR Package provides the basic functions to deal with each step of the pipeline. Finally, a step-by-step guide for installing and utilizing the framework is reported at https://qbioturin.github.io/connector/.

To make accessible the basic CONNECTOR functionalities even to users without expertise in R, we developed a web application through the R package Shiny (RStudio, Inc, 2014), a powerful R library for developing interactive web apps straight from R. Specifically, the function *RunConnectorShiny* offers a fancy web application providing a basic-level interface to configure and execute a data analysis with the CONNECTOR functionalities. Thus, users without experience in R language can directly focus on analyzing the results rather than spend their efforts setting up an R script. Therefore, the web application enables the user to run each function and go through the three steps of analysis with just a few clicks. All these functionalities make the CONNECTOR Package a very flexible and easy tool.

### Xenograft In Vivo Treatments

After surgical removal from patients, each metastatic colorectal cancer specimen was fragmented and either frozen or prepared for implantation: cut in small pieces of which 2 fragments were implanted in 2 mice. After engraftment and tumor mass formation, the tumors were passaged and expanded for 2 generations until production of 2 cohorts, each consisting of 12 mice. Tumor size was evaluated once-weekly by calliper measurements and the approximate volume of the mass was calculated using the formula 4/3*π*(*d*/2)^2^*D*/2, where *d* is the minor tumor axis and *D* is the major tumor axis. PDXs derived from each original fragment were then randomized for treatment with placebo (6 mice) or cetuximab (6 mice); animals with established tumors, defined as an average volume of 400*mm*^3^, were then treated with cetuximab (Merck, White House Station, NJ) 20 mg/kg/twice-weekly i.p.

For assessing PDX models response to therapy, we used averaged volume measurements at 3 weeks after treatment normalized to the tumorgraft volume at the time of cetuximab treatment initiation. Tumors are then classified as follows:

1. “partial response” (**PR**): decrease of at least 50% in tumor volume
2. “progressive disease” (**PD**): increase of at least a 35% in tumor volume
3. “stable disease” (**SD**): the ones between 50% decrease and 35% increase

All animal procedures were approved by the Ethical Commission of the Institute for Cancer Research and Treatment and by the Italian Ministry of Health.

### Transcriptional analyses

Differentially expressed genes (DEGs) between different CTGC or keratin-high and -low groups were obtained using the R package DESeq2 (v1.26.0) (Love *et al*., 2014) with design: batch+cluster. Here batch is used to correct for sequencing batches and cluster to select samples belonging to the CTGC of interest (or keratin-high and -low). We did not compare CTGC clusters for which less than 5 samples were available. Genes with more than 5 reads in only 1 sample were removed before testing for differential expression, DEGs were identified using |LFC| >=0.5849625 and adjusted p-value < 0.05. Gene Ontology analyses for the upregulated or downregulated genes were performed with the R library ClusterProfiler (v3.14.3) (Wu *et al*., 2021; Yu *et al*., 2012).

The selection of a robust set of markers for the epithelial stratification differentiation phenotype was performed with two criteria:

- being assigned to the GO keratinization (GO:0031424 (ebi, 2022)), the most significantly enriched term across all the CTGC-Ac vs other CTGC comparisons, via direct experimental evidence, using filters for taxon (Homo Sapiens) and evidence (manual assertion) on the EBI Quick GO web site. **AND** being significantly upregulated either in Bb versus Ac or Bc versus Ac

The resulting genes are: KRT80, KRT7, KRT86, KRT81, KRT83, KRT6B, KRT6A, KRT74, KRT79, KRT16. To classify samples in high or low expressing keratin groups, quartiles of the expression level of the ten selected genes were calculated from progressive disease (PD) samples belonging to CTGC-Ac. Each sample was then assigned ten labels, according to its expression of those keratins with respect to CTGC-Ac first and third quartiles - “high” if the expression is larger than the 3rd quartile and “low” if it is smaller than the 1st. Samples were then classified as “keratin-high” if they were assigned no low labels, and “keratin-low” if they were assigned no high labels.

Z transformed expression values in COAD/READ patients for HOPX were downloaded alongside clinical annotations from CBioPortal (Firehose legacy dataset), patients with multiple expression values or missing expression/survival data were filtered out. Survival analysis was performed on the remaining 371 patients using the libraries survival and survminer (Therneau, 2022; Kassambara, 2021), the optimal threshold for HOPX expression was determined using the maximally selected rank statistic approach (maxstat, (Hothorn, 2017)), correcting accordingly the resulting log rank p value. DEG and all the enrichment and survival analyses were performed with R version 3.6.3.

### Morphological analyses

Haematoxilin and Eosin was performed in xenografts from mice treated with vehicle (until tumors reached an average volume of 1500*mm*^3^). Tumors were explanted, routinely processed and stained with H&E (Bio-optica). Images were captured with the Leica LAS EZ software using a Leica DM LB microscope.

### Further computational methods

#### Naive Bayesian Classification

The *naive Bayesian classification* is a probabilistic classifier based on the Bayes’ theorem. It can be efficiently exploited to assign a cluster membership to each parental tumor (from now on referred as model) given the cluster membership of every curve of the PDXs derived from it. Let 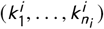 be the clusters assigned to the *n_i_*, PDX curves belonging to the *i*th model. Thus, the classifier assigns the cluster 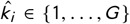 to the *i*th model as:

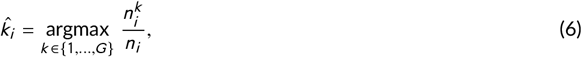

where 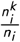 is the frequency of having 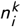 curves of the *i*th model in the *k*th cluster.

#### Shannon Index

The *Shannon index* (Shannon, 1948) is an information statistic index that we used for quantifying the curve diversity in a specific PDX model. It is computed as follows:

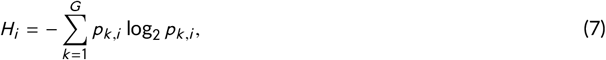

where *H_i_*, is the Shannon index for the *i*th model, *G* is the number of clusters, *p_k,i_*, is the proportion of PDX curves from the *i*th model and belonging to the *k*th cluster. Therefore, a larger value of *H_i_*, corresponds to a larger diversity of the *i*th PDX model, i.e. many different clusters are assigned to the same model. Whether *H_i_*, is equal to zero means that the curves of the *i*th PDX model belong to the same cluster.

#### t-distributed stochastic neighbor embedding (t-SNE)

t-SNE (Van der Maaten and Hinton, 2008) is a statistical method used to visualize high-dimensional data in a two or three-dimensional map exploiting a nonlinear dimensionality reduction technique. This approach has been applied to give a spatial visualization of the resulting CTGCs of the PDS models in Figure 3C.

## Acknowledgements

We would like to thank Eugenia Zanella for xenografts cetuximab experiments, Massimiliano Frassà for working on the LAS and Caterina Marchiò for a second blind evaluation of the H&E sections. We would like also to express our thanks to Jude Fitzgibbon, Marco Ladetto and Simone Ferrero for the useful discussions, and to Jessica Giordano and Elena Rosso for the critical check of CONNECTOR.

## Author contributions

Concept and design (FC, RS, SP, and MO); Develop of CONNECTOR (SP, RS, and MB); Data analyses and interpretation (SP, RS, EG, AB, and FC); Provisions of study materials for OV case study (MO, LA, DT, and RAC); Provisions of study materials for mCRC case study (LT,AB, EG, MV, MF, and FS); manuscript writing (SP, RS, AB, EG, and FC); manuscript editing (SP, RS, EG, MV, MF, FS, LA, DT, RAC, LV, AB, MB, MO, and FC); all authors gave final approval of the version.

## Supporting Information

CONNECTORwebsite: https://qbioturin.github.io/connector Protocols.io link: dx.doi.org/10.17504/protocols.io.8epv56e74g1b/v1.

## Supporting Information

**FIGURE S1.**
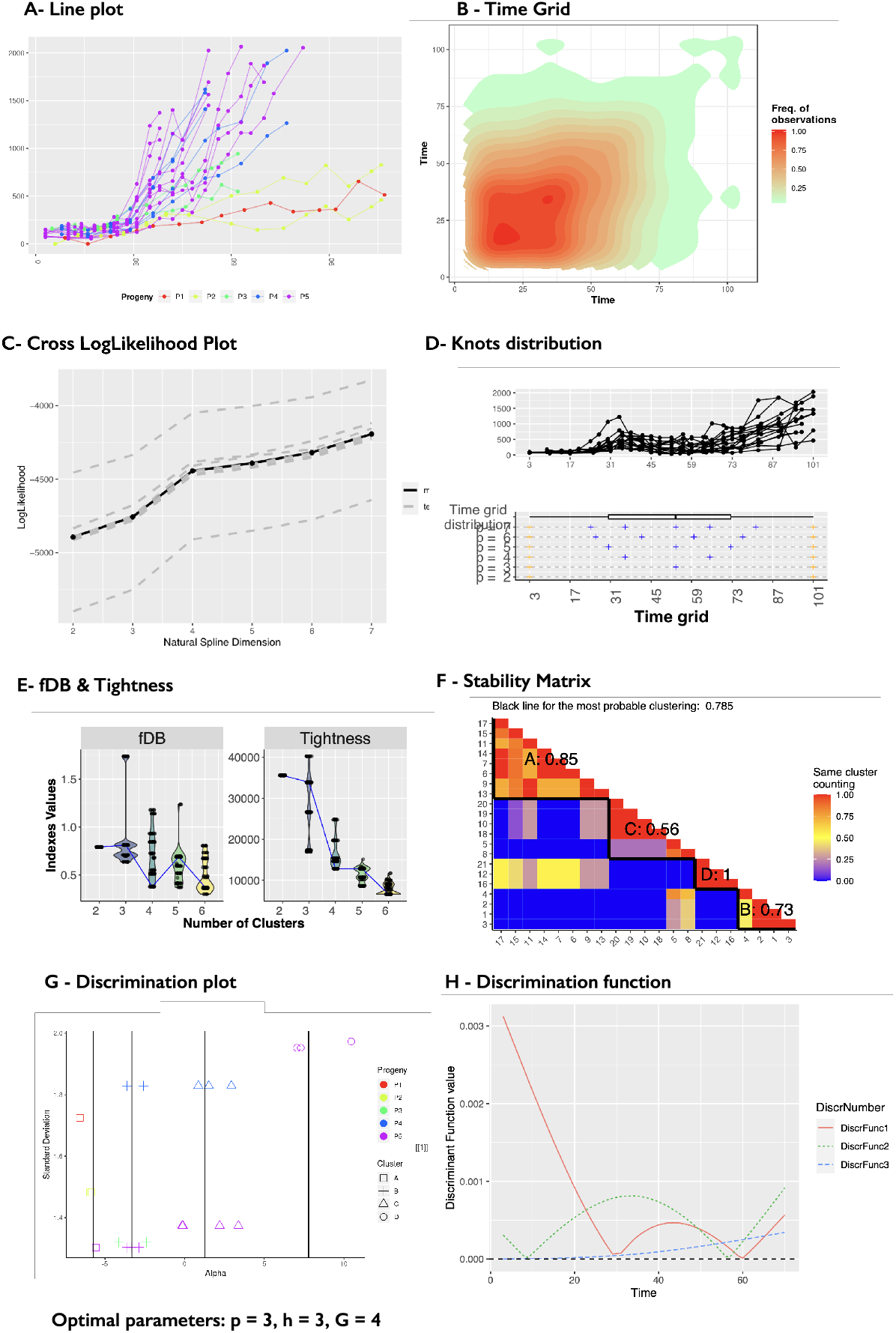
**A** line plot of the 21 PDX curves of HGS-EOC tumor. **B** Time grid, the suggested time at which truncate the data is 70 days. **C** Cross LogLikelihood plot reports 4 as the optimal parameter for the base spline, this value is also supported by the Knots distribution, **D**. **E** fDB and tightness violin plot for the selection of the number of clusters, fours is associated with the lower fDB value and at 4 is is appreciable the elbow in the tightness plot. **F** the stability matrix obtained selecting 4 clusters. **G** Discriminant plot and **H** discriminant functions. The optimal set of parameters selected is p=3, h=3, and G=4.

**FIGURE S2.**
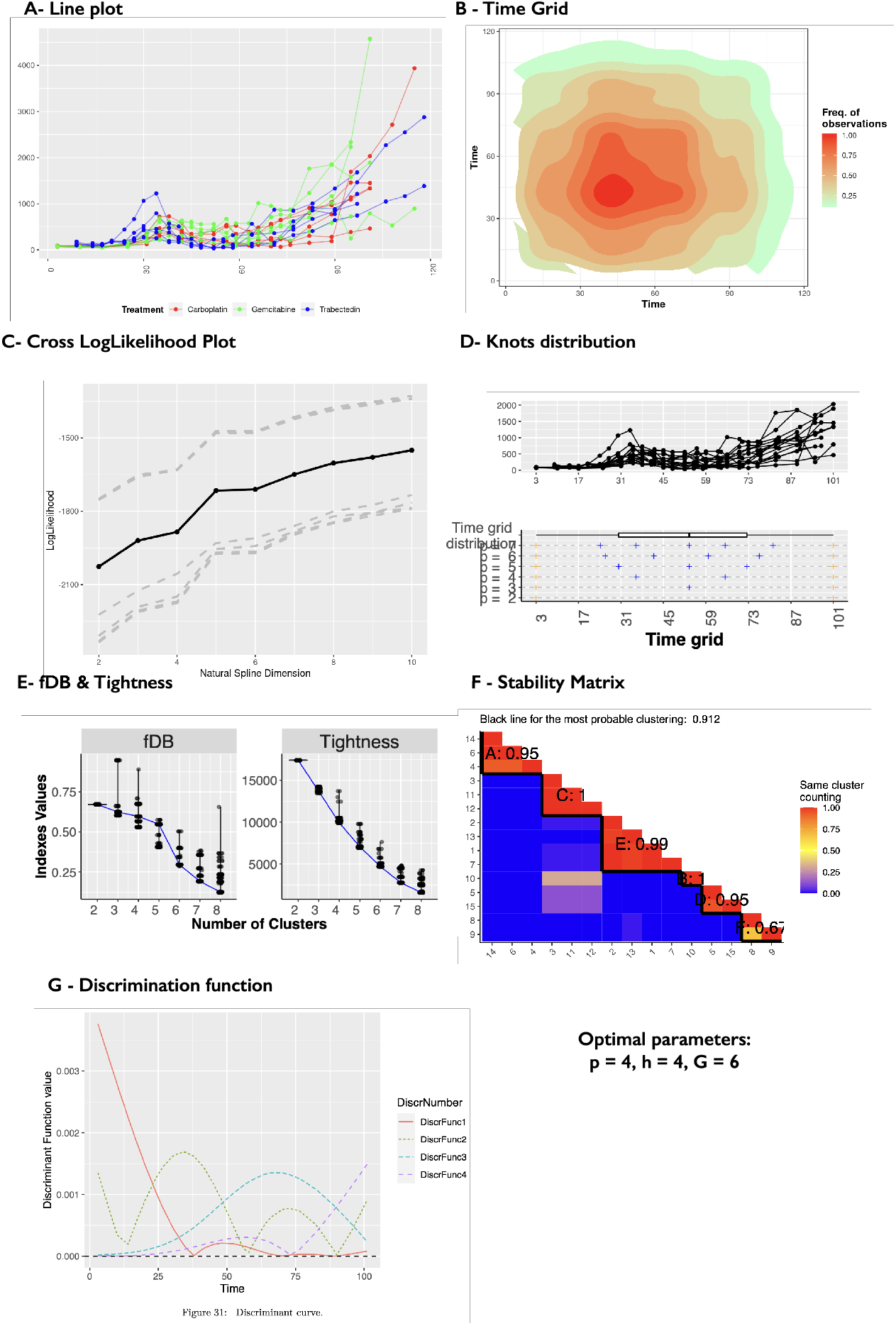
**A** line plot of the 25 PDX curves of HGS-EOC tumor treated with three drugs treatments, carboplatin, gemcitabine and trabectedin, reported in red, green and blue, respectively. **B** Time grid, the suggested time at which truncate the data is 70 days. **C** Cross LogLikelihood plot reports 4 as the optimal parameter for the base spline, this value is also supported by the Knots distribution, **D. E** fDB and tightness violin plot for the selection of the number of clusters, six is associated with the lower fDB value. At 6 is appreciable a slightly elbow in the tightness plot. **F** the stability matrix obtained selecting 6 clusters. **G** the discriminant functions are reported. The optimal set of parameters selected is p=4, h=4, and G=6.

**FIGURE S3.**
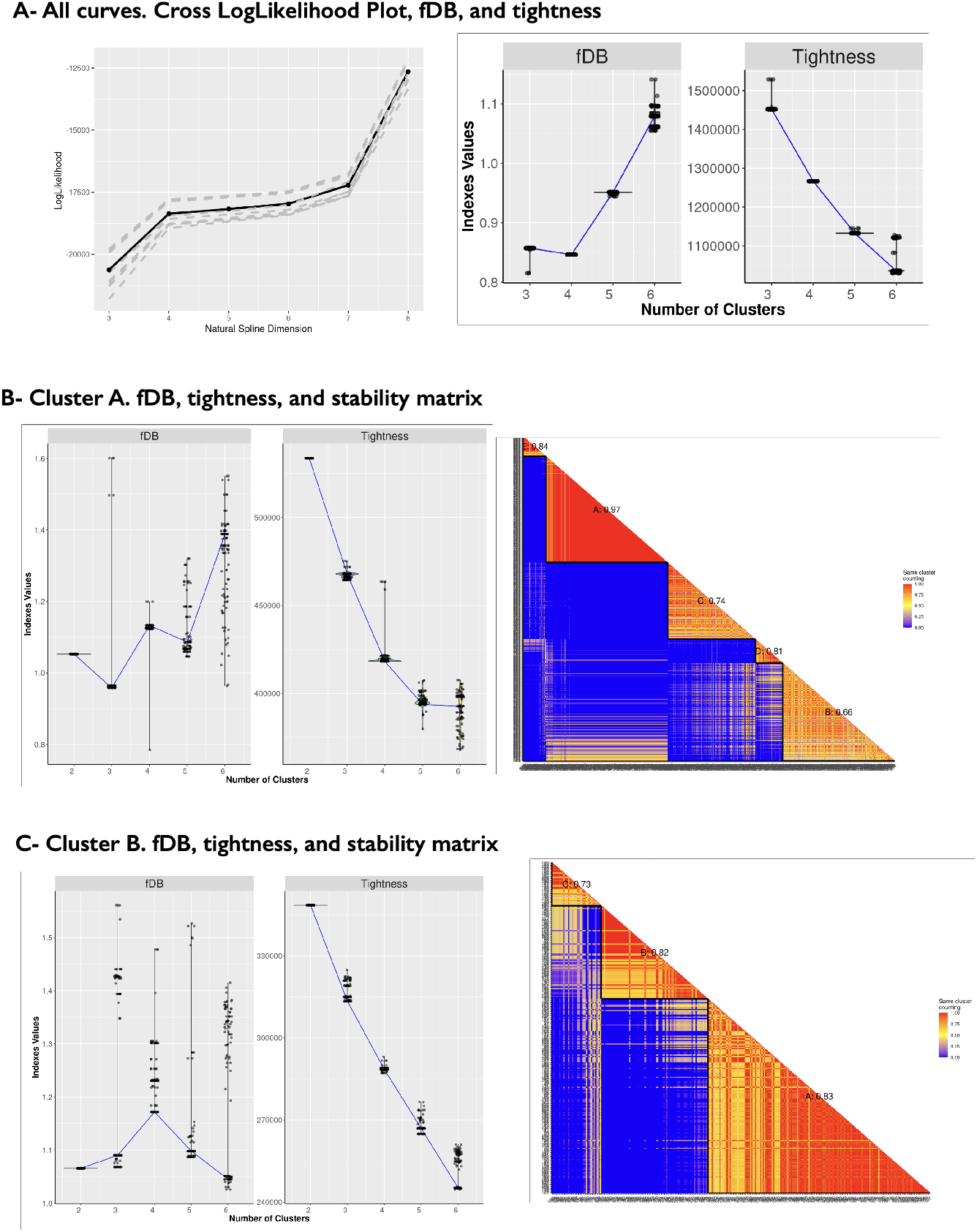
**A** Cross LogLikelihood plot, fDB and tightness obtained from the CONNECTOR run on all 1536 individual mice. The optimal set of parameters selected is p=4, h=2, and G=3. **B** fDB, tightness grid, and stability matrix obtained on cluster A. The optimal set of parameters selected is p=4, h=3, and G=5. **C** fDB, tightness grid, and stability matrix obtained on cluster B. The optimal set of parameters selected is p=4, h=3, and G=3.

**FIGURE S4.**
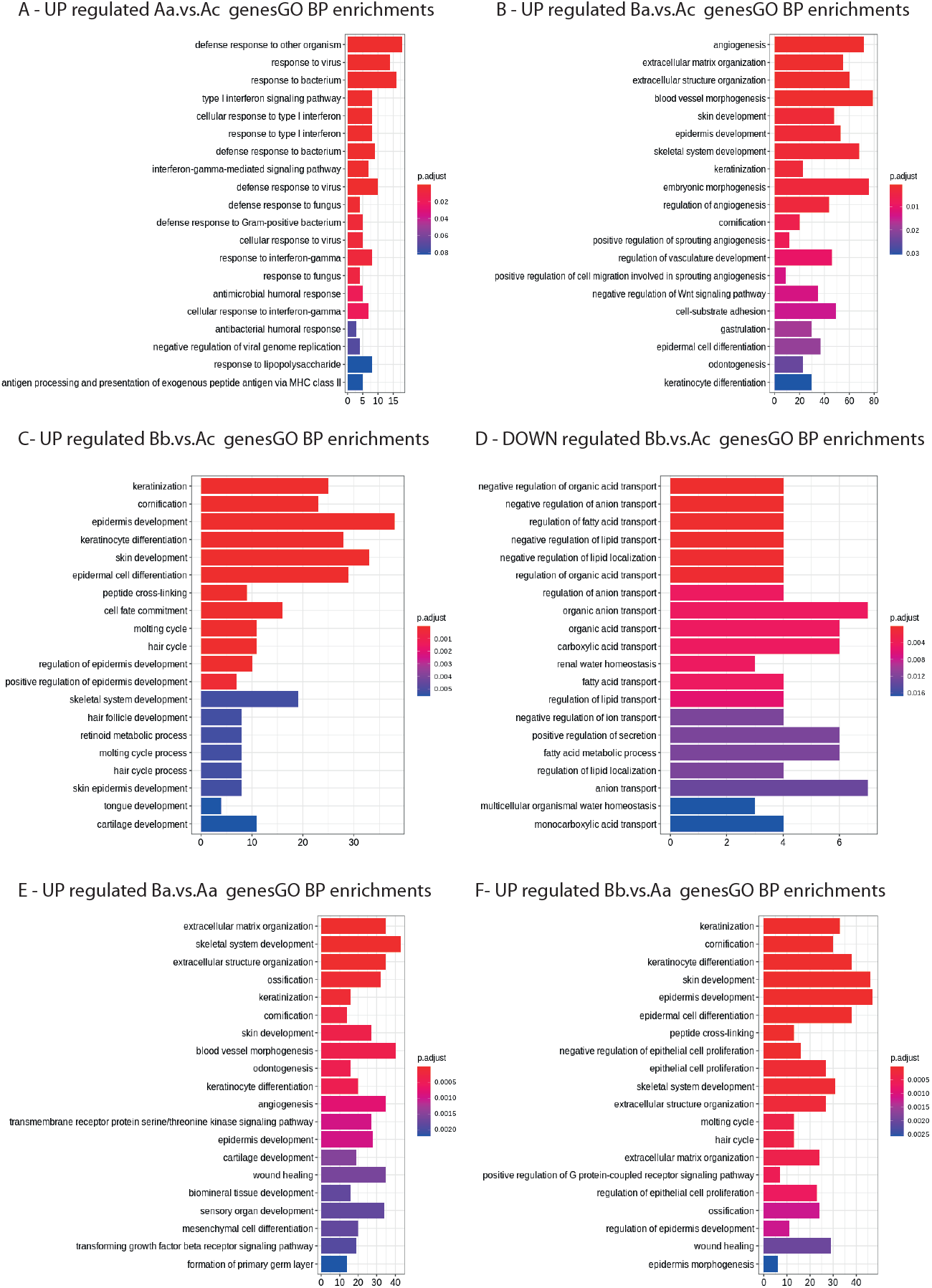
GO BP enrichments for DEG comparing CTGC: Only comparisons that yielded statistically significant (BH adjusted p value <0.001) terms are shown. Specifically A CTGC-Aa vs CTGC-Ac; B CTGC-Ba vs CTGC-Ac; C and D CTGC-Bb vs CTGC-Ac; E CTGC-Ba vs CTGC-Aa and F CTGC-Bb vs CTGC-Aa. Height of the bars indicate the number of genes in the GO and color shading the adjusted p value.

**FIGURE S5.**
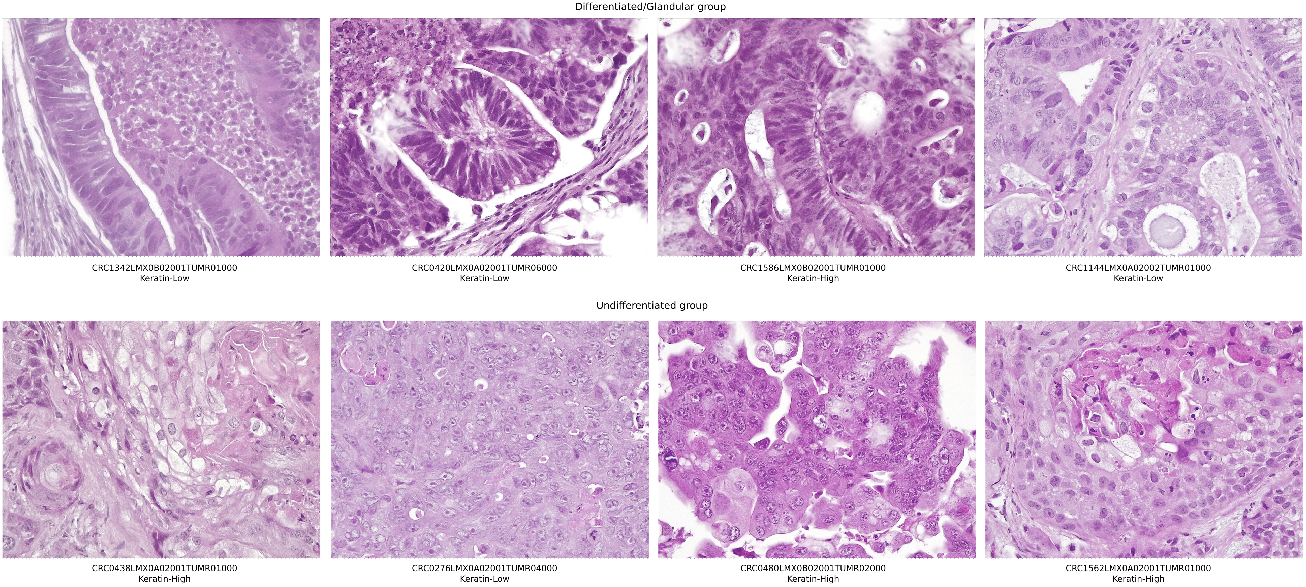
H& morphological analysis clearly shows two distinct groups: Haematoxilin and Eosin (H&E) was performed in blind on eight tumors from the two groups. On the top tumors are displaying histological features of a more differentiated phenotype, with clear adenomorphic structures. On the bottom, tumors are less differentiated and less structured. keratin-high and -low annotations are also shown.

**FIGURE S6.**
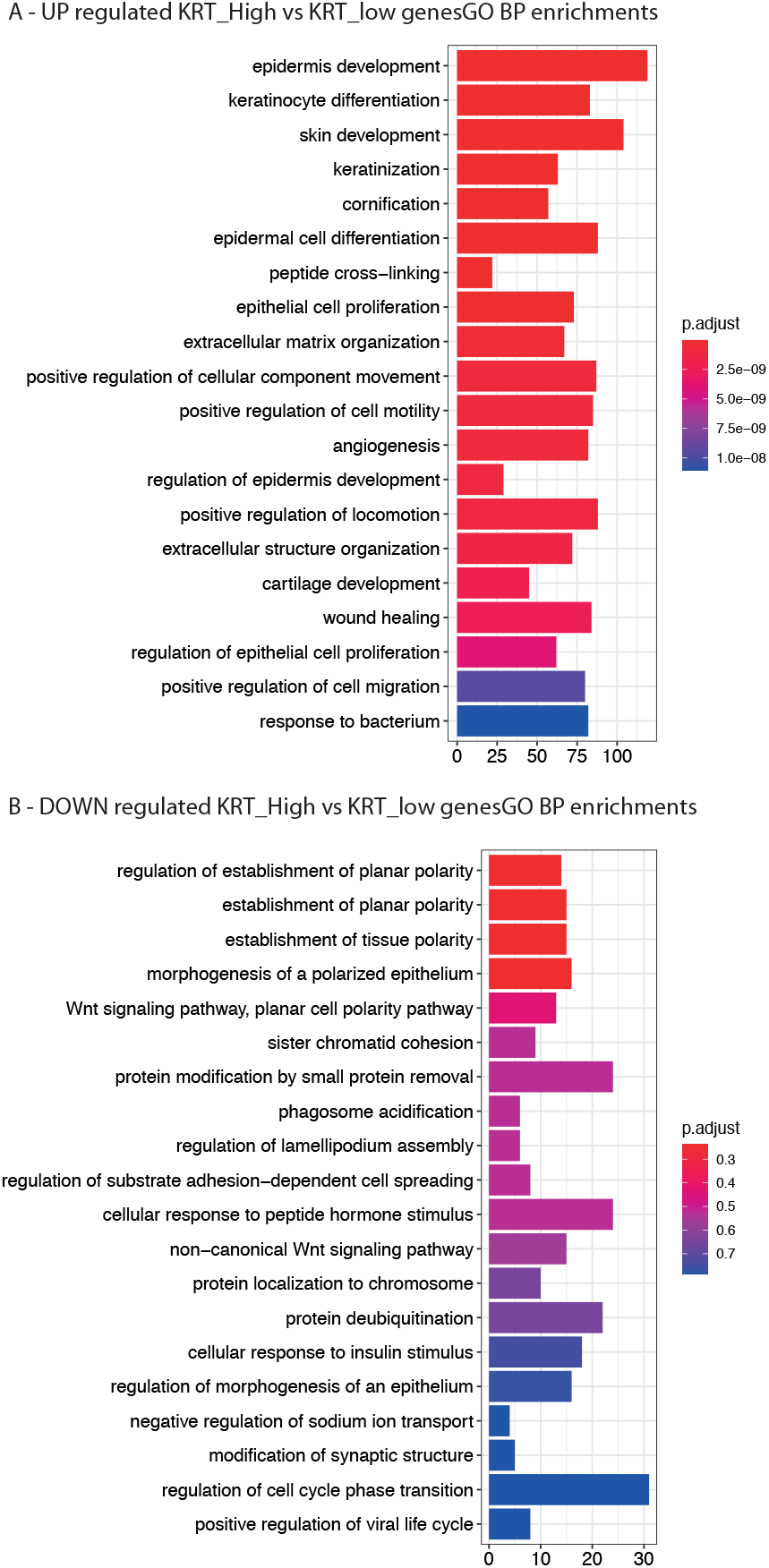
GO BP enrichments for DEG between keratin-high and keratin-low tumors: Barplots of Gene Ontology results based on the DEG between 42 keratin-high and 15 keratin-low samples. Both upregulated (A) and downregulated (B) signatures were represented. Height of the bars indicate the number of genes in the GO and color shading the adjusted p value. Downregulated genes are not associated with any significant signature.

### MODEL SELECTION TOOLS: NUMERICAL DETAILS

The parametrized family of semi-metrics between curves, denoted as *D_q_* and defined in eq. (3), is the building block for both the *total tightness* T defined in eq. (4) and the *functional* DB (fDB) defined in eq. (5). Such indexes are proposed as measures for the quality of the clustering of functional data, useful for a proper comparison between different classifications and as a guide to the user. Let’s here illustrate the numerical details needed for their reliable calculation.

Following Ferraty and Vieu (2006b), the numerical evaluation of *D_q_* should exploit the spline approximation of the curves involved. For *q* = 0 the curves are not differentiated and hence eqs. (S2) and (S1) give the values of the curves at any points. Hence, the integral in eq. (3) can be evaluated using an appropriate quadrature formula on the full time grid, made of all the sampled time points for all the sampled curves, as CONNECTOR calculates curve predictions and cluster means for the entire time interval.

For *q* ≥ 1, the computation of successive derivatives is performed by direct calculation of their analytical form. More precisely, the estimated *i*-th curve is given by eq. (S2) as a linear combination of the spline basis functions **s**(*t*) (with known analytical expressions) and the coefficients 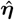

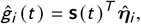

and the differentiated estimated *i*–th curve can be represented

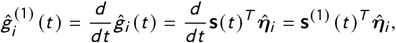

as where the analytical expressions of **s**^(1)^(*t*) are known too, being the derivatives of the basis functions **s**(*t*). Let us comment that, the functional clustering method James and Sugar (2003) consider the spline basis **s**(*t*) obtained by the singular value decomposition of the B-spline basis for a natural cubic spline. Hence the (*k, p*) matrix *A* with elements the values of the B-spline basis (dimension *p*) functions evaluated at the grid time points (*k*) is decomposed

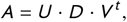

as where *U* and *V* are (*k, k*) and (*p, p*) orthogonal matrices and *D* is a (*k, p*) diagonal with non zero elements (*p*_1_,…, *p_p_*). The spline basis considered in James and Sugar (2003) is given by the first *p* columns of the matrix *U* (the only columns which enter the product with matrix *D*). Hence for *i* = 1,…, *p*, the *i*-th colum of such matrix is given by the linear transformation

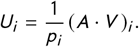

When *q* ≥ 1, the same linear transformation applies to the matrix *B* with elements the *q*-th derivative of the natural B-spline basis functions evaluated at the grid time points. So finally the integral in definition (3) can be evaluated with a suitable quadrature formula.

### S2 FUNCTIONAL CLUSTERING MODEL: DETAILS

Notice that **λ**_0_, Λ and ***α**_k_* are confounded if no constraints are imposed. Hence, we ask

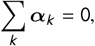

meaning that **s**(*t*)^T^**λ**_0_ may be interpreted as the overall mean curve. Moreover, we ask

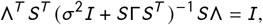

the reason of which is well explained in (James and Sugar, 2003).

Notice that the *k*th cluster mean curve can be retrieved as

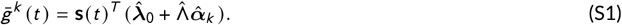

Moreover, the functional clustering procedure can accurately predict unobserved portions of the curves *g_i_*(*t*) by means of the natural estimate

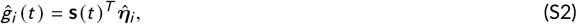

where 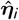, is a prediction for **η**_*i*_, which is proven to be optimally computed as 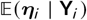 and explicitly given in (James and Sugar, 2003), eq. (17).

### S3 PERFORMANCE OF DIFFERENT CLUSTERING METHODS ON SPARSE AND IRREGULARLY SAMPLED CURVES

In this Section we review the performances of different clustering procedures that could be implemented on longitudinal data.

#### Clustering with Classical Growth Models

As we address growth curves in this paper, we cannot skip to mention the classical growth models that have been introduced in the literature. In the following, we review the main growth models (Malthus, Gompertz and logistic) and the performance of a clustering method based on the direct estimation of the parameters involved. Indeed, each sampled curve could be represented as the array of the estimated values of the parameters of the fitted growth model. In doing so, the full sample of curves becomes a sample of points in 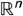, where *n* is the number of parameters of the growth model, and the clustering procedure can be performed using classical clustering methods for data points.

Let us briefly review the most popular growth models. These models are usually formulated in terms of ordinary differential equations (ODEs) that relate the growth rate of the tumour to its current state and range from the simple one-parameter exponential growth model to more advanced models that contain more parameters.

The *Malthus model* was one of the first mathematical models introduced to study and model exponential-linear population growth. It assumes that every cell continuously passes through the cell cycle giving birth to two daughter cells at regular intervals. Then, the number of cancer cells and therefore the volume of the tumour would increase exponentially over time. In particular, the Malthus model is described by the following ODE:

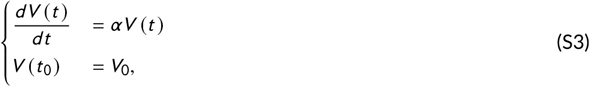

where *V*(*t*) is the total tumour volume at time *t*, *α* is a constant for the growth rate, and *V*_0_ is the tumour volume at the initial time *t*_0_.

Successively, the *Gompertz model* was derived from a generalisation of the Malthus model, assuming that the relative growth rate is not constant, but time dependent with a decreasing behaviour at constant positive rate *β*:

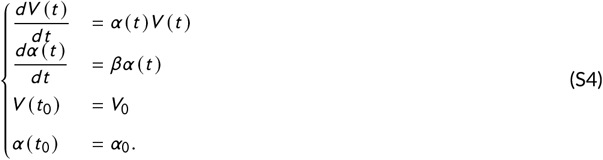

Finally, in the *logistic model*, the relative growth rate decreases linearly, when the population size increases, according with the assumption that resources are limited. Thus, the ODEs defining the model are:

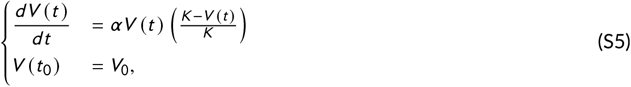

where *α* is a coefficient related to population kinetic and *K* is the so-called carrying capacity, which represents the maximum population size a particular environment can support.

For each model above reported, a clustering algorithm can be implemented in a two steps procedure:

- For every *i*, estimate the model parameters by fitting the *i*-th sampled curve. The result is an array (*p*_*i*_1__,…, *p*_*i_*n*_*_), where *n* is the number of model parameters. Each sampled curve is now represented as a data point in dimension *n*.
- Cluster the full sample of data points 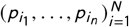 by a classical clustering algorithm. We chose a k-means algorithm.

The procedure has been tested and the performance is reported here.

##### The comparison setup

We tested the clustering with classical growth models on four sets of growth curves, see Figure S7. The four CONNECTOR analyses are reported at https://qbioturin.github.io/connector/examples/.

For each sample, we first fitted the Malthus, Gompertz and logistic models. The estimated parameters for each curve in each of the four samples are reported in Tables S4, S5, S6 and S7.

The data points are then clustered through a k-means clustering algorithm. The optimal number of clusters is determined by evaluating the *Total Tightness*, see eq. (4) (in the main paper), and the *fDB indexes*, see eq. (5) (in the main paper), calculated using the distance between curves, see eq. (3) (in the main paper). The quality of the final clustering of the corresponding curves is evaluated by the fDB index. The results are reported in Table S8. Those values will be compared to the values of the same index on clustering of the same samples through functional methods, discussed in the following paragraph. As a first comment, notice that the classical models are not flexible enough to fit the curves in test 4, which exhibit more complex dynamics than a “simple” growth.

**FIGURE S7.**
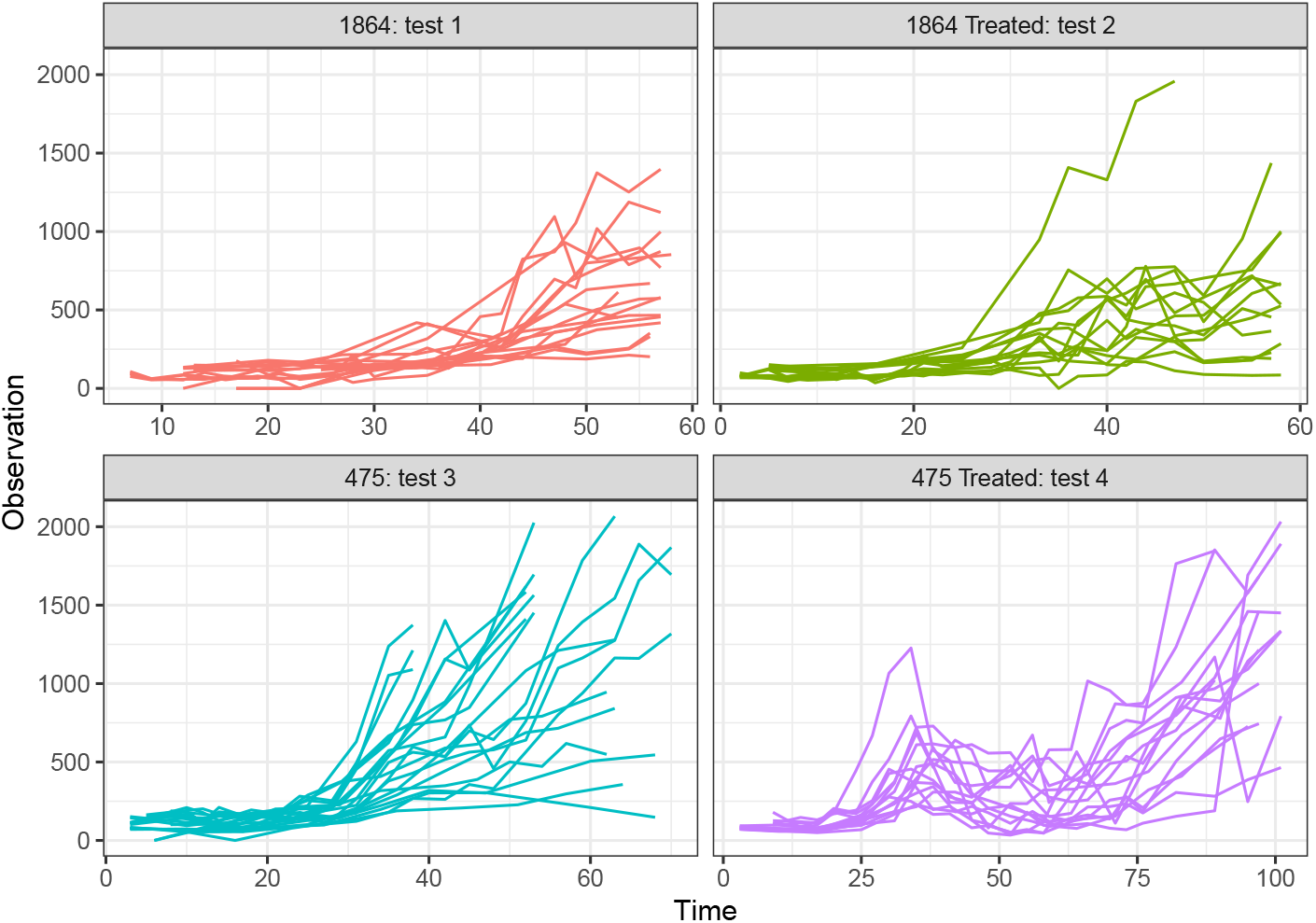
The four tests samples.

#### Review and Comparison of Functional Clustering Methods

Clustering functional data is generally a difficult task because of the infinite dimensional space from which data are sampled. Different approaches have been proposed along the years. Two nice reviews on functional data clustering can be found here (Müller, 2005), (Jacques and Preda, 2014).

The most popular approach consists of reducing the problem to a finite dimensional setting by approximating data with elements from some finite dimensional space. Afterwards, clustering algorithms for finite dimensional data can be run. The reducing dimension step, often denoted as *filtering* step, consists in approximating the curves into a finite basis of functions. Spline basis is one of the most common choice because of their optimal properties. Another dimension reduction technique is the functional principal component analysis, based on the Karhunen-Loeve expansion of a square integrable *L*^2^ stochastic process. The implementation of functional principal components also requires some form of regularisation, which can be achieved with smoothing methods, see (Ramsay and Silverman, 2005).

On the other hand, nonparametric methods for clustering have been proposed, see (Ferraty and Vieu, 2006a). They consist generally in defining specific distances or dissimilarities for functional data and then apply clustering algorithms as hierarchical clustering or k-means.

Finally, model-based clustering techniques have been developed. In this cases the observations are modeled as mixture distributions. Two main currents have been explored: one which models the functional principal components scores (Bouveyron and Jacques, 2011) and another one which models directly the expansion coefficients in a finite basis of functions (James and Sugar, 2003).

The advantages and disadvantages of each method are summarized in (Jacques and Preda, 2014). We do not enter the discussion here and refer the reader to the original paper. As the conclusion of the critical analysis presented in (Jacques and Preda, 2014) is that model-based techniques are better performing, we decided to rely on such techniques. In particular on the two methods presented in (James and Sugar, 2003) and in (Bouveyron and Jacques, 2011). We tested both techniques on the “general growth curve” type of data which are the object of this paper.

**Table S4.** Estimated parameters for each sampled curve in Test 1.

| Malthus |  | Logistic |  |  | Gompertz |  |  |
| --- | --- | --- | --- | --- | --- | --- | --- |
| $V_0$ | $\alpha$ | $V_0$ | $K$ | $\alpha$ | $V_0$ | $\alpha_0$ | $\beta$ |
| 68.78308 | 0.04445 | 39.39801 | 1701.15668 | 0.06589 | 31.56854 | 0.08716 | 0.01548 |
| 36.89524 | 0.04989 | 6.40114 | 656.23237 | 0.10991 | 1.83531 | 0.24387 | 0.03739 |
| 11.61218 | 0.07304 | 11.55314 | 99999.99076 | 0.07323 | 11.61144 | 0.07304 | 0 |
| 28.55079 | 0.05322 | 15.54528 | 1221.30806 | 0.07456 | 17.80942 | 0.07694 | 0.00828 |
| 22.0811 | 0.05481 | 1.35357 | 499.86858 | 0.14483 | 0.00012 | 1.08476 | 0.07014 |
| 20.43031 | 0.0543 | 4.40454 | 531.67804 | 0.10741 | 0.08896 | 0.44936 | 0.05004 |
| 40.12871 | 0.0477 | 39.95205 | 99999.91164 | 0.04787 | 40.12695 | 0.0477 | 0 |
| 91.71518 | 0.0414 | 0.62068 | 889.00398 | 0.19828 | 0 | 1.85772 | 0.0893 |
| 47.13489 | 0.05348 | 46.75451 | 99999.80804 | 0.05378 | 47.13253 | 0.05348 | 0 |
| 125.2202 | 0.00929 | 84.84368 | 210.67975 | 0.06516 | 78.17013 | 0.05303 | 0.05259 |
| 90.93223 | 0.01975 | 90.91487 | 99999.99218 | 0.01979 | 90.93134 | 0.01975 | 0 |
| 46.32061 | 0.04917 | 20.80527 | 1135.41884 | 0.07941 | 11.54091 | 0.13244 | 0.02368 |
| 65.85038 | 0.02732 | 65.81126 | 99999.8859 | 0.02737 | 65.84926 | 0.02732 | 0 |
| 25.90587 | 0.06918 | 0.0866 | 1198.44381 | 0.21987 | 0 | 1.36521 | 0.06464 |
| 18.81232 | 0.0585 | 13.72665 | 1920.26111 | 0.06913 | 18.81129 | 0.0585 | 0 |
| 27.59306 | 0.07141 | 0.51379 | 1598.25243 | 0.17844 | 0 | 1.39001 | 0.06416 |
| 22.62525 | 0.06707 | 0.17156 | 1014.09897 | 0.19457 | 2e-05 | 1.10764 | 0.05973 |

##### The comparison setup

The two selected clustering procedures are both implemented in a software. The clustering procedure introduced in (Bouveyron and Jacques, 2011) is implemented in the **R** package ‘funHDDC, maintained and available for download from CRAN. The method by (James and Sugar, 2003) is implemented in an **R** function ‘fclust’ available directly from James’s webpage *http://faculty.marshall.usc.edu/gareth-james/index.html*.

We tested both methods on four sets of growth curves from Figure S7, the same dataset on which the clustering with classical growth models have been tested, see Table S8. The results are summarised in Table S9. Notice that ‘funHDDC’ includes a model selection function to help the user set the optimal number of clusters, while ‘fclust’ has been integrated with the model selection procedures introduced in CONNECTOR. Moreover, the functional latent mixture model presented in (Bouveyron and Jacques, 2011) can be reduced to different submodels by constraining model parameters within or between groups. Here we tested the submodels which are denoted as [*a_kj_, b_k_*, *Q_k_*, *d_k_*], [*a_k_, b_k_, Q_k_, d_k_*], [*a_kj_, b, Q_k_, d_k_*] and [*a_k_, b, Q_k_, d_k_*].

Notice that both fclust and funHDDC result in very good fDB indexes, compared to the fDB indexes reported in Table S8 for the classical models. Both the functional clustering methods are based on very flexible models, that can adapt to data. On the contrary, the simple classical models have too few degrees of freedom and cannot adjust to the sampled curves.

**Table S5.**
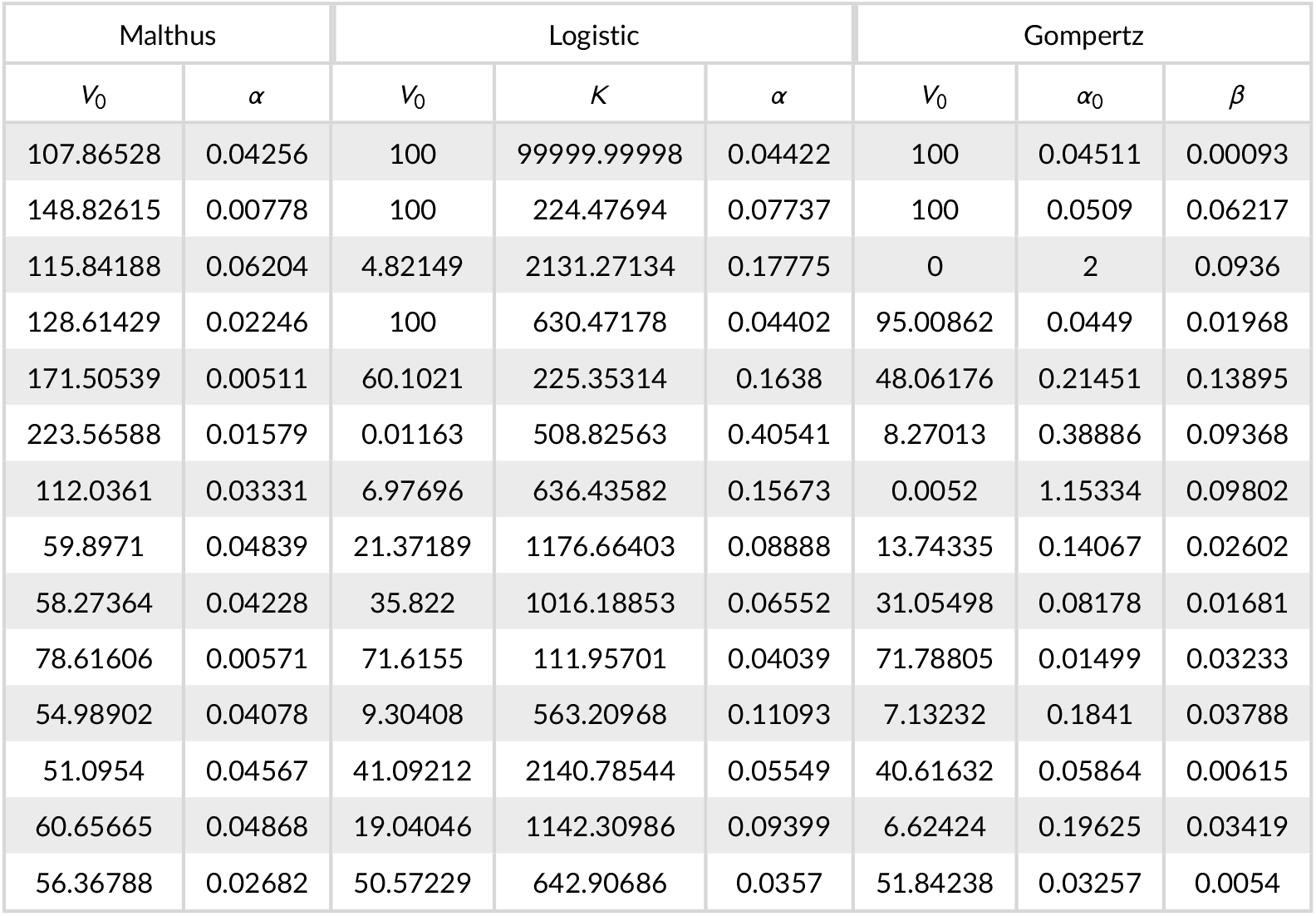
Estimated parameters for each sampled curve in Test 2.

As shown in Table S9, the method by (James and Sugar, 2003) lead to far better results. The conclusion of the comparison integrates the study performed in (Jacques and Preda, 2014), as here, strongly sparse and irregularly sampled curves are considered, few points per curve ~ 7 to 20 instead of ~ 31 to 241.

**Table S6.**
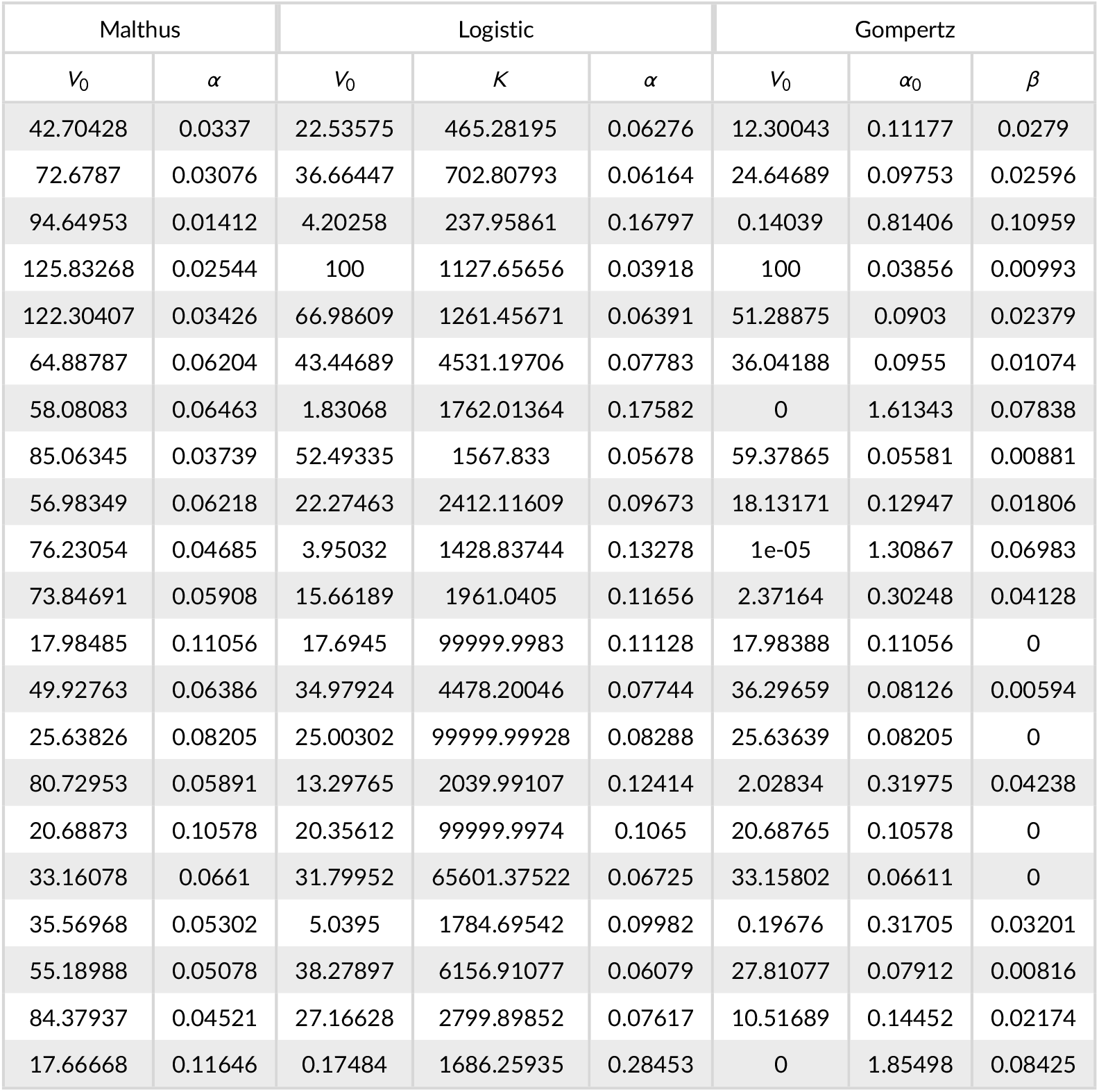
Estimated parameters for each sampled curve in Test 3.

| Malthus |  | Logistic |  |  | Gompertz |  |  |
| --- | --- | --- | --- | --- | --- | --- | --- |
| $V_0$ | $\alpha$ | $V_0$ | $K$ | $\alpha$ | $V_0$ | $\alpha_0$ | $\beta$ |
| 42.70428 | 0.0337 | 22.53575 | 465.28195 | 0.06276 | 12.30043 | 0.11177 | 0.0279 |
| 72.6787 | 0.03076 | 36.66447 | 702.80793 | 0.06164 | 24.64689 | 0.09753 | 0.02596 |
| 94.64953 | 0.01412 | 4.20258 | 237.95861 | 0.16797 | 0.14039 | 0.81406 | 0.10959 |
| 125.83268 | 0.02544 | 100 | 1127.65656 | 0.03918 | 100 | 0.03856 | 0.00993 |
| 122.30407 | 0.03426 | 66.98609 | 1261.45671 | 0.06391 | 51.28875 | 0.0903 | 0.02379 |
| 64.88787 | 0.06204 | 43.44689 | 4531.19706 | 0.07783 | 36.04188 | 0.0955 | 0.01074 |
| 58.08083 | 0.06463 | 1.83068 | 1762.01364 | 0.17582 | 0 | 1.61343 | 0.07838 |
| 85.06345 | 0.03739 | 52.49335 | 1567.833 | 0.05678 | 59.37865 | 0.05581 | 0.00881 |
| 56.98349 | 0.06218 | 22.27463 | 2412.11609 | 0.09673 | 18.13171 | 0.12947 | 0.01806 |
| 76.23054 | 0.04685 | 3.95032 | 1428.83744 | 0.13278 | 1e-05 | 1.30867 | 0.06983 |
| 73.84691 | 0.05908 | 15.66189 | 1961.0405 | 0.11656 | 2.37164 | 0.30248 | 0.04128 |
| 17.98485 | 0.11056 | 17.6945 | 99999.9983 | 0.11128 | 17.98388 | 0.11056 | 0 |
| 49.92763 | 0.06386 | 34.97924 | 4478.20046 | 0.07744 | 36.29659 | 0.08126 | 0.00594 |
| 25.63826 | 0.08205 | 25.00302 | 99999.99928 | 0.08288 | 25.63639 | 0.08205 | 0 |
| 80.72953 | 0.05891 | 13.29765 | 2039.99107 | 0.12414 | 2.02834 | 0.31975 | 0.04238 |
| 20.68873 | 0.10578 | 20.35612 | 99999.9974 | 0.1065 | 20.68765 | 0.10578 | 0 |
| 33.16078 | 0.0661 | 31.79952 | 65601.37522 | 0.06725 | 33.15802 | 0.06611 | 0 |
| 35.56968 | 0.05302 | 5.0395 | 1784.69542 | 0.09982 | 0.19676 | 0.31705 | 0.03201 |
| 55.18988 | 0.05078 | 38.27897 | 6156.91077 | 0.06079 | 27.81077 | 0.07912 | 0.00816 |
| 84.37937 | 0.04521 | 27.16628 | 2799.89852 | 0.07617 | 10.51689 | 0.14452 | 0.02174 |
| 17.66668 | 0.11646 | 0.17484 | 1686.25935 | 0.28453 | 0 | 1.85498 | 0.08425 |

**Table S7.**
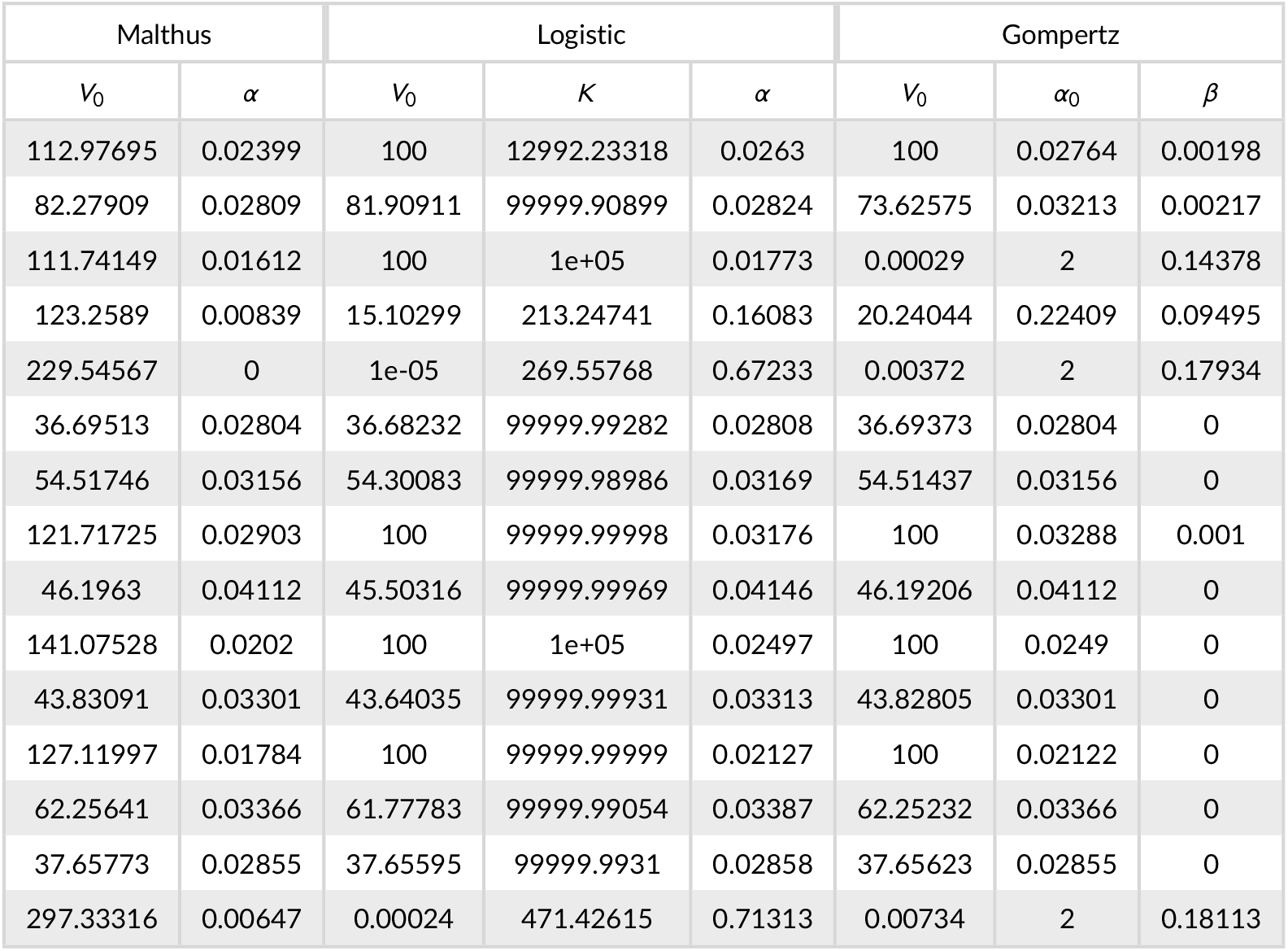
Estimated parameters for each sampled curve in Test 4.

| Malthus |  | Logistic |  |  | Gompertz |  |  |
| --- | --- | --- | --- | --- | --- | --- | --- |
| $V_0$ | $\alpha$ | $V_0$ | $K$ | $\alpha$ | $V_0$ | $\alpha_0$ | $\beta$ |
| 112.97695 | 0.02399 | 100 | 12992.23318 | 0.0263 | 100 | 0.02764 | 0.00198 |
| 82.27909 | 0.02809 | 81.90911 | 99999.90899 | 0.02824 | 73.62575 | 0.03213 | 0.00217 |
| 111.74149 | 0.01612 | 100 | 1e+05 | 0.01773 | 0.00029 | 2 | 0.14378 |
| 123.2589 | 0.00839 | 15.10299 | 213.24741 | 0.16083 | 20.24044 | 0.22409 | 0.09495 |
| 229.54567 | 0 | 1e-05 | 269.55768 | 0.67233 | 0.00372 | 2 | 0.17934 |
| 36.69513 | 0.02804 | 36.68232 | 99999.99282 | 0.02808 | 36.69373 | 0.02804 | 0 |
| 54.51746 | 0.03156 | 54.30083 | 99999.98986 | 0.03169 | 54.51437 | 0.03156 | 0 |
| 121.71725 | 0.02903 | 100 | 99999.99998 | 0.03176 | 100 | 0.03288 | 0.001 |
| 46.1963 | 0.04112 | 45.50316 | 99999.99969 | 0.04146 | 46.19206 | 0.04112 | 0 |
| 141.07528 | 0.0202 | 100 | 1e+05 | 0.02497 | 100 | 0.0249 | 0 |
| 43.83091 | 0.03301 | 43.64035 | 99999.99931 | 0.03313 | 43.82805 | 0.03301 | 0 |
| 127.11997 | 0.01784 | 100 | 99999.99999 | 0.02127 | 100 | 0.02122 | 0 |
| 62.25641 | 0.03366 | 61.77783 | 99999.99054 | 0.03387 | 62.25232 | 0.03366 | 0 |
| 37.65773 | 0.02855 | 37.65595 | 99999.9931 | 0.02858 | 37.65623 | 0.02855 | 0 |
| 297.33316 | 0.00647 | 0.00024 | 471.42615 | 0.71313 | 0.00734 | 2 | 0.18113 |

**Table S8.** Test results for the classical models.

|  | Malthus |  | Gompertz |  | logistic |  |
| --- | --- | --- | --- | --- | --- | --- |
|  | Number of clusters | fDB index | Number of clusters | fDB index | Number of clusters | fDB index |
| test 1 | $G = 4$ | 3.753 | $G = 4$ | 1.9956 | $G = 5$ | 4.8957 |
| test 2 | $G = 5$ | 0.8628 | $G = 6$ | 2.1597 | $G = 5$ | 1.3979 |
| test 3 | $G = 5$ | 1.631 | $G = 4$ | 0.8986 | $G = 6$ | 0.7092 |
| test 4 | $G = 6$ | 1.7203 | $G = 3$ | 2.0086 | $G = 4$ | 2.0892 |

**Table S9.** Test results from fclust and funHDDC.

|  | fclust |  |  | funHDDC |  |  |
| --- | --- | --- | --- | --- | --- | --- |
|  | Parameters | Number of clusters | fDB index | Best Model Parameters | Number of clusters | fDB index |
| test 1 | $p = 3$<br>$h = 1$ | $G = 4$ | 0.4831 | $[a_k, b, Q_k, d_k]$<br>$p = 3$ | $G = 2$ | 0.7932 |
| test 2 | $p = 3$<br>$h = 1$ | $G = 4$ | 0.4594 | $[a_k, b, Q_k, d_k]$<br>$p = 4$ | $G = 3$ | 0.8853 |
| test 3 | $p = 3$<br>$h = 1$ | $G = 4$ | 0.3897 | $[a_k, b, Q_k, d_k]$<br>$p = 4$ | $G = 3$ | 0.5539 |
| test 4 | $p = 3$<br>$h = 2$ | $G = 4$ | 0.4049 | $[a_k, b, Q_k, d_k]$<br>$p = 3$ | $G = 3$ | 1.4478 |

